# A Multiplexed SEC-MS Approach to Systematically Study the Interplay Between Protein Assembly-States and Phosphorylation Events

**DOI:** 10.1101/2023.01.12.523793

**Authors:** Ella Doron-Mandel, Benjamin J. Bokor, Yanzhe Ma, Lena A. Street, Lauren C. Tang, Ahmed A. Abdou, Neel H. Shah, George A. Rosenberger, Marko Jovanovic

## Abstract

A protein’s molecular interactions and post-translational modifications (PTMs), such as phosphorylation, can be co-dependent and reciprocally co-regulate each other. Although this interplay is central for many biological processes, a systematic method to simultaneously study assembly-states and PTMs from the same sample is critically missing. Here, we introduce SEC-MX (Size Exclusion Chromatography fractions MultipleXed), a global quantitative method combining Size Exclusion Chromatography and PTM-enrichment for simultaneous characterization of PTMs and assembly-states. SEC-MX enhances throughput, allows phosphopeptide enrichment, and facilitates quantitative differential comparisons between biological conditions. Applying SEC-MX to HEK293 and HCT116 cells, we generated a proof-of-concept dataset mapping thousands of phosphopeptides and their assembly-states. Our analysis revealed intricate relationships between phosphorylation events and assembly-states and generated testable hypotheses for follow-up studies. Overall, we establish SEC-MX as a valuable tool for exploring protein functions and regulation beyond abundance changes.

## Introduction

A major effort in studying the dynamics of biological systems is to measure differential changes in the proteome. Classically, proteome dynamics have been studied by measuring differences in total protein expression levels. However, total proteome levels do not convey the full array of post-translational dynamics. For example, the protein’s assembly-state – whether a protein is acting alone as a monomer or as part of different complexes – often changes between biological conditions and systems, despite similarities in overall expression levels^1–3^. These differences in assembly-states, often driven by protein-protein interactions (PPIs), may reflect distinct functions that contribute to differential cell states.

An additional central mechanism contributing to the complexity of the proteome is the attachment of functional groups to proteins, referred to as post-translational modifications (PTMs)^4^. Notable examples encompass ubiquitination, acetylation, methylation, and phosphorylation – which is one of the most extensively studied. PTMs can induce various alterations in protein activity, including activation or inhibition, tagging for degradation, and subcellular localization, among others. Notably, most PTMs and PPIs are dynamic and take part in regulatory mechanisms, and therefore change between biological conditions^5–9^. Therefore, characterizing these dynamic “states” is essential for a comprehensive biological understanding.

Cumulative research over several decades has demonstrated the interdependence of a protein’s assembly-state and PTM-status^5,8–11^. For instance, phosphorylation can modulate a protein’s interaction interface, and conversely, protein interactors can obstruct phosphorylation sites, thereby denying access to kinases or phosphatases. Past works on the interplay between PPIs and PTMs have relied on targeted methods, such as co-immunopurification, to enrich the interactomes of specific proteoforms^5,8^. However, these approaches are limited in their scope and a global scale method to simultaneously study assembly-states and PTMs is critically missing. Therefore, we set out to develop an approach to systematically study the interplay between protein assembly-states and phosphorylation events.

Previously, Size Exclusion Chromatography followed by Mass Spectrometry (SEC-MS) has been used successfully to analyze assembly-states, identify PPIs and characterize the composition of molecular-complexes^1,12–26^. SEC is used to distinguish different assembly-states of the same protein, which elute in distinct fractions based on the different molecular weight (MW) of each assembly (for example, a monomer will elute in a low MW fraction versus a multimeric complex of a higher MW^2^). However, SEC-MS is limited by input amounts that render PTM enrichment challenging. As a result, previous works have mined phosphopeptides in SEC datasets by meta-analysis^27^, or studied the effects of phosphatase-treatment on SEC elution profiles^28^. However, no study to date has directly measured enriched phosphopeptides with SEC-MS.

Here, we introduce a global and quantitative methodology that enables the simultaneous characterization of PTMs and assembly-states, in the same sample, to explore their relationship across various biological processes. Expanding on prior SEC-MS and co-fractionation multiplexing works^29^, we use isobaric tags to develop SEC-MX – Size Exclusion Chromatography fractions MultipleXed – and demonstrate its advantages in; (1) improving throughput by reducing the number of LC-MS/MS runs required to reconstruct the PPI network, (2) enabling phosphopeptide enrichment and measurements thereby allowing characterization of PTMs along the SEC range, and (3) simplifying quantitative comparisons between biological conditions that are multiplexed together.

In this study, we employed SEC-MX to comprehensively characterize the SEC elution profiles of both non-modified and phosphorylated peptides from HEK293 and HCT116 cells. This yielded a novel dataset, marking the first instance of concurrent non-targeted measurements of phosphorylation events and assembly-states for thousands of proteins across two distinct biological conditions. Our analysis enabled a comparative examination of assembly-states and their phosphorylation status between the two cell-lines. Overall, our study provides insights into the intricate interplay between post-translational modifications and protein assembly-states, underscoring the unique value of SEC-MX in unraveling the complexities of protein regulation.

## Results

### Development and benchmarking of SEC-MX

We developed SEC-MX (SEC fractions MultipleXed) to enable quantitative comparison between different samples, as well as to measure phosphorylation events on proteins in distinct assembly-states by enriching phosphopeptides from SEC fractions. First, we set out to multiplex SEC fractions to increase the sample-yield for phosphopeptide enrichment. Multiplexing was achieved by labeling SEC fractions with isobaric tags. Tandem mass tags (TMTpro) were used because they provide the highest number of labeling channels currently available, allowing combinations of up to 18 samples in a single liquid-chromatography tandem mass spectrometry (LC-MS/MS) run. We multiplexed SEC-adjacent fractions within the same TMT mix to minimize the occurrence of missing data points and reduce sample complexity (Figure 1A-B, Extended Data Figure 1A). In doing so, we took advantage of the fact that once a peptide is triggered for MS2 acquisition, TMT reporter intensity values are, in most cases, assigned to all channels and therefore achieve more complete elution profiles. In addition, we designed a “full-overlap” scheme (Figure 1B, Extended Data Figure 1C) in which every fraction is measured twice, in different mixes, increasing coverage and enabling batch correction between different TMT-mixes (the development of the mixing scheme is detailed in the Materials and Methods section, Extended Data Figure 1).

**Figure 1.**
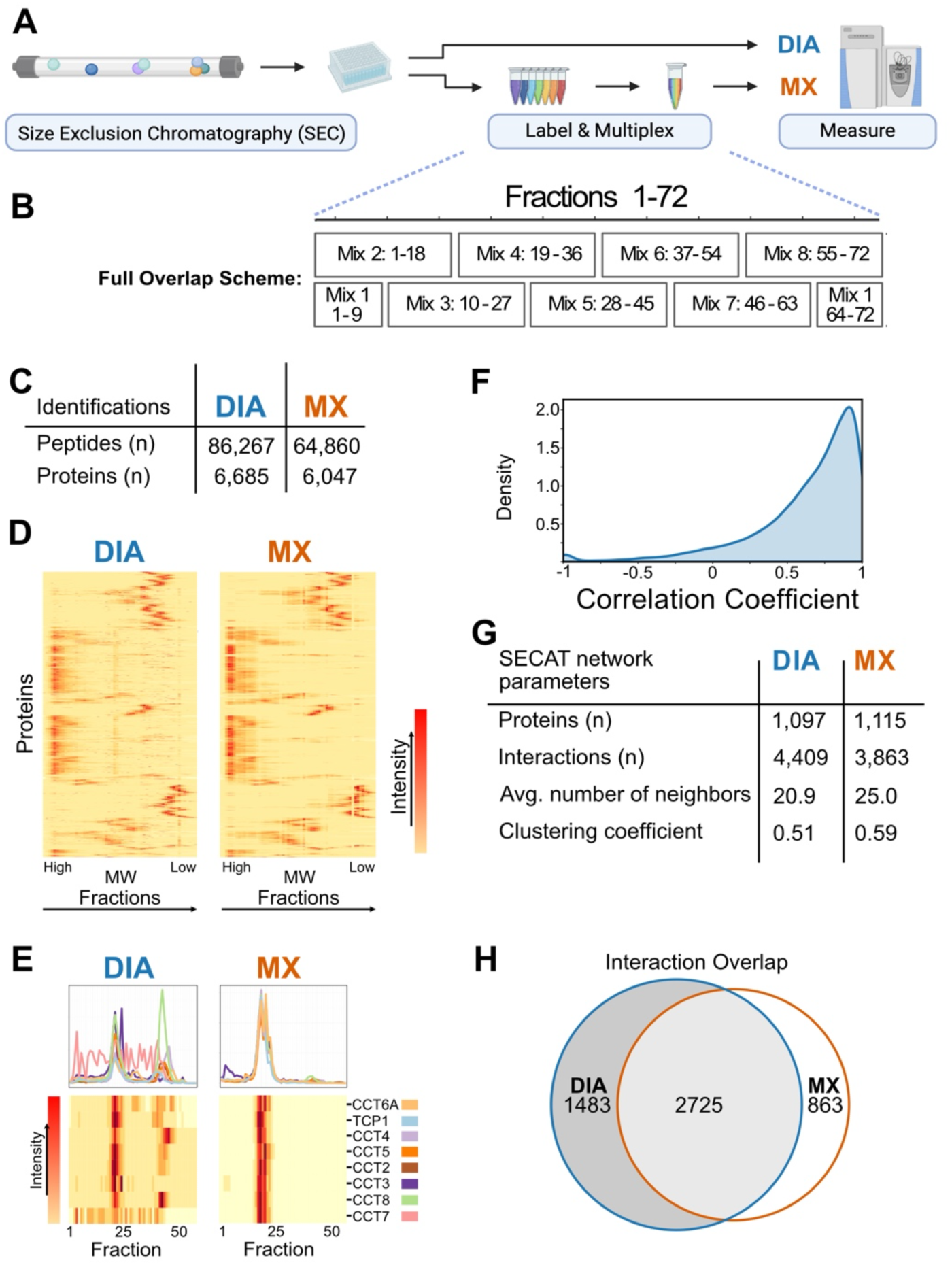
SEC-MX performs comparably to SEC-DIA in coverage and resolution: ***(A)*** Overview of experimental pipeline: Cells are lysed under physiological conditions, followed by fractionation on a size exclusion chromatography (SEC) column into ∼90 fractions. The protein-containing fractions (54-72 fractions total) are further processed by tryptic digestion. The resulting peptides are either measured individually using DIA (SEC-DIA), or labeled by TMT, multiplexed and measured in pools of 18 fractions (SEC-MX). **(B)** A “full overlap” mixing scheme was developed, in which each fraction is divided in two and each half is measured in a different mix, keeping adjacent fractions together. **(C)** Protein and peptide identifications in SEC-DIA and SEC-MX. **(D)** Heatmap representation comparing signals in SEC-DIA and SEC-MX, for proteins measured in both. Columns represent fractions, rows represent different proteins, which are row-normalized from 0 to 1 so that the max elution peak per protein is represented in red. Rows in both heatmaps are arranged in the same order. **(E)** Elution traces and heatmap representation of the SEC elution of the CCT complex subunits in either SEC-DIA or SEC-MX. **(F)** Distribution of Pearson correlation coefficients between elution profiles in SEC-MX versus SEC-DIA per protein measured in both. **(G)** Parameters of the PPI networks build by SECAT analysis of either SEC-DIA or SEC-MX data. **(H)** The overlap of interactions between SEC-DIA and SEC-MX, q-value < 0.05 in at least one condition and < 0.1 in the other, (see Materials and Methods section for details).

To compare SEC-MX to the field’s current gold standard – label-free SEC measured with data independent acquisition (DIA), we conducted SEC on HEK293 cells and measured the resulting fractions either individually or multiplexed by TMT labeling, in duplicate biological replicates for each. We found that coverage in SEC-MX was comparable to SEC-DIA, with only a 10% difference in protein-group identifications, despite a 9-fold reduction in the number of LC-MS/MS runs (57 versus 8 in DIA or TMT, respectively) (Figure 1C). Additionally, SEC-MX produced very similar SEC elution patterns to that of SEC-DIA, evident by the positions of the elution peaks per protein, as well as in the high-correlation between elution profiles measured by both methods (Figure 1D-F). We then used the network-centric analysis algorithm SECAT ^22^ to identify high-confidence PPIs and observed that SEC-MX identified a similar number of PPIs as did SEC-DIA, (3,588 and 4,208 at 5% false-discovery rate, respectively, of which 54% were overlapping), with similar network parameters (Figure 1G-H, Extended Data Figure 2A-B). We additionally analyzed the data with a reference-free PPI analysis tool which allows identifications of novel interactions^24^, resulting in 12,620 and 14,406 PPIs in SEC-MX and SEC-DIA, respectively (Extended Data Figure 2C-E).

Together, our observations showed that SEC-MX performs comparably to the field’s gold standards in building a context-specific human PPI network while requiring an order of magnitude less LC-MS/MS runs.

### SEC-MX enables phospho-peptide enrichment

After validating SEC-MX performance for studying protein interactions, we next set out to test the feasibility of the multiplexing approach in enabling phosphopeptide enrichment using standard immobilized metal affinity chromatography (IMAC). As mentioned above, we were interested in developing the method in order to facilitate differential analysis between biological conditions. Therefore, we designed the study to compare between two distinct cell-lines, HEK293 and HCT116. Fifty-four SEC fractions were collected from each cell-line, in two biological replicates. To minimize variability in protein coverage and to increase the chances of identifying the same phosphopeptides from the two cell-lines, we multiplexed fractions from both samples together in the same TMT mixes (Figure 2A, Extended Data Figure 1B). After labeling and multiplexing, 20% of each mix were taken for measuring the ‘global’ proteome for analysis of protein assembly-states, while the rest of the sample was further allocated for phosphopeptide enrichment using IMAC (Figure 2A).

**Figure 2.**
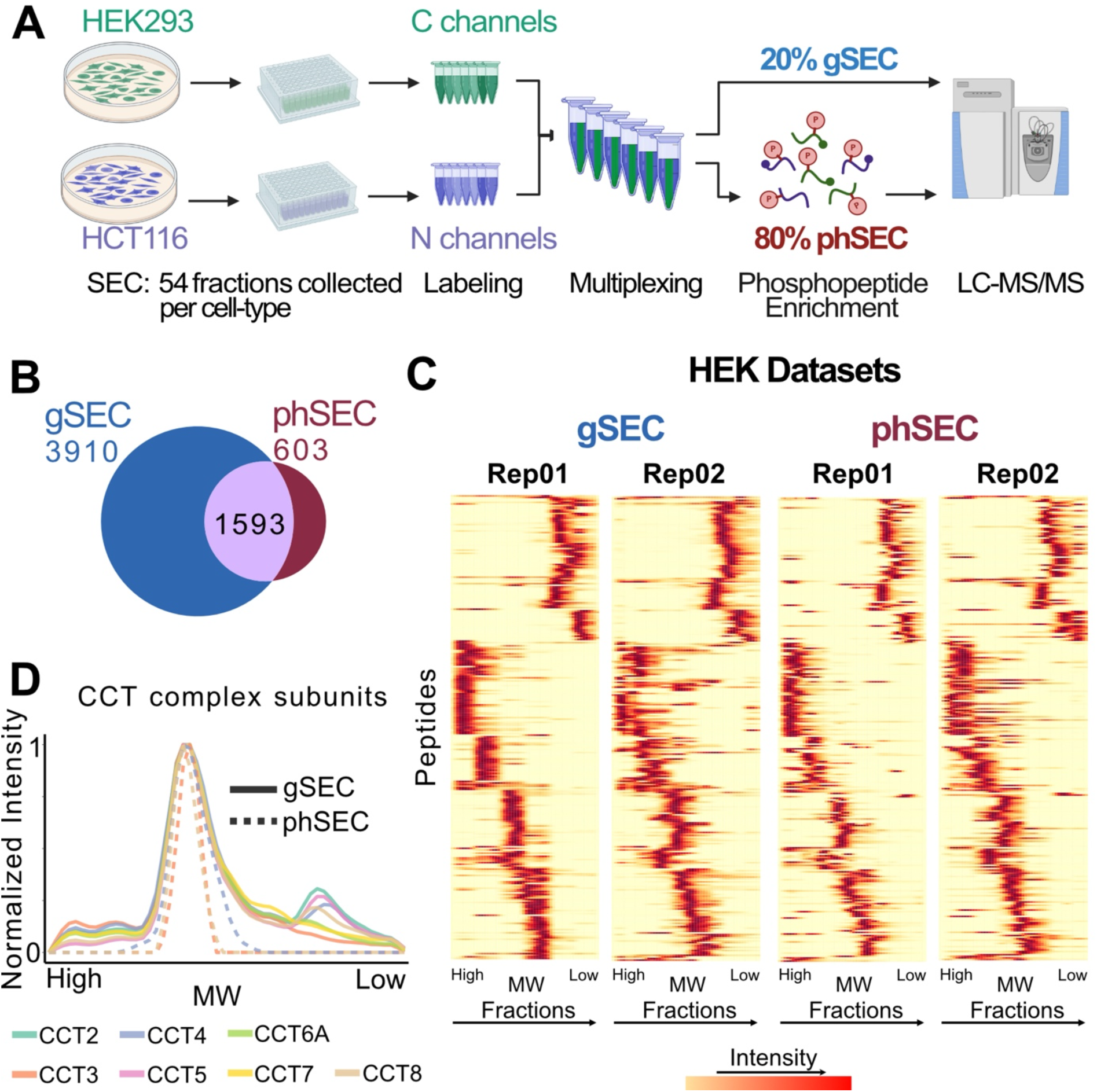
SEC-MX enables phosphopeptide enrichment: **(A)** Overview of experimental setup. The same fractions from HEK293 or HCT116 cells were multiplexed together, digested, and labeled. After pooling, 80% of the sample was allocated for phosphopeptide enrichment and measurement of phSEC, while the remaining 20% was processed for gSEC. **(B)** Venn diagram showing the overlap of protein identifications between the gSEC and phSEC datasets. **(C)** Heatmap representation of the elution profiles of overlapping peptides in gSEC and phSEC from HEK293 cells in both replicates. Columns represent fractions, rows represent different peptides, which are scaled from 0 to 1 so that the max elution peak per protein is represented in red. Rows in all 4 heatmaps are arranged in the same order. For similar heatmaps from HCT116 cells see Extended Data Figure 3. **(D)** Elution traces for CCT complex members identified in gSEC and phSEC (HEK293 cells, average of both replicates, similar results were obtained for HCT116 cells, shown in Extended Data Figure 3).

Overall, the global dataset (hereafter referred as gSEC) yielded 59,659 peptides, covering 5,503 protein groups (Table 1). In the phosphopeptide SEC dataset (phSEC) we recovered 4,762 phosphorylated peptides, spanning 2,196 proteins, out of which 1,593 overlapped with the gSEC dataset (Figure 2B). To initially assess the quality of the phSEC data, we compared the elution patterns of peptides in phSEC to their corresponding peptides in gSEC and observed a high degree of correlation (Figure 2C, Extended Data Figure 3A-D). For example, we inspected the elution profiles of phosphopeptides mapping to subunits of the stable CCT complex and found that they co-eluted with the fully assembled form of the complex (Figure 2D, Extended Data Figure 3E). Of note, since the information content of different phosphopeptides is not necessarily linked to each other, meaning they might not belong to the same proteoforms, from here on we report all phSEC data on the peptide-level and match it to the protein-level data of the parent-protein in the gSEC dataset (3,609 phosphopeptides, matched to 1,593 proteins). In conclusion, our PTM enrichment downstream from SEC fractionation created a unique and first-of-its-kind dataset measuring SEC elution profiles for ∼4000 phosphopeptides and their matching parent proteins, which enables overlaying PTM-status with protein assembly-states.

**Table 1.** Overview of assembly-state relevant identifications per dataset.

| Groups |  | Numbers |  |  |  |  |  |
| --- | --- | --- | --- | --- | --- | --- | --- |
| Replicate | Condition | Dataset | Proteins | Peptides | Pep Peaks | Prot Peaks | Assembly States |
| 01 | HCT | gSEC | 4,578 | 41,429 | 57,234 | 7,747 | 8,519 |
|  |  | phSEC | 2,027 | 4,237 | 6,314 |  |  |
|  | HEK | gSEC | 4,583 | 41,466 | 52,236 | 7,083 |  |
|  |  | phSEC | 1,993 | 4,159 | 5,473 |  |  |
| 02 | HCT | gSEC | 5,154 | 46,763 | 69,702 | 9,508 | 11,592 |
|  |  | phSEC | 1,477 | 2,745 | 4,101 |  |  |
|  | HEK | gSEC | 5,151 | 46,743 | 71,240 | 10,005 |  |
|  |  | phSEC | 1,483 | 2,761 | 3,837 |  |  |

### A protein-centric analysis framework to study assembly-states

Next, we aimed to analyze our unique dataset in order to; (1) define assembly-states for every measured protein, (2) map phosphorylation events onto assembly-states and, (3) study how these patterns differ between conditions. Previously, SEC-MS has been widely used to identify pair-wise PPIs, characterize the composition of molecular-complexes, or compare interactions between samples^1,18,22,24,30^. As opposed to a focus on interactions, recent work has shown the potential of a protein-centric approach that looks at the assembly-state changes of individual proteins between conditions ^2^. Based on this approach, we first identified assembly-states by taking into account that each SEC elution peak represents a distinct assembly-state of the protein (Figure 3A). This allowed us to assign/map phosphorylation events by aligning individual peaks between gSEC and phSEC. Lastly, we used a protein-centric approach to compare those assembly-states between biological conditions and discover potential regulatory events that occur on specific assembly-states of the same protein (Figure 3A).

**Figure 3.**
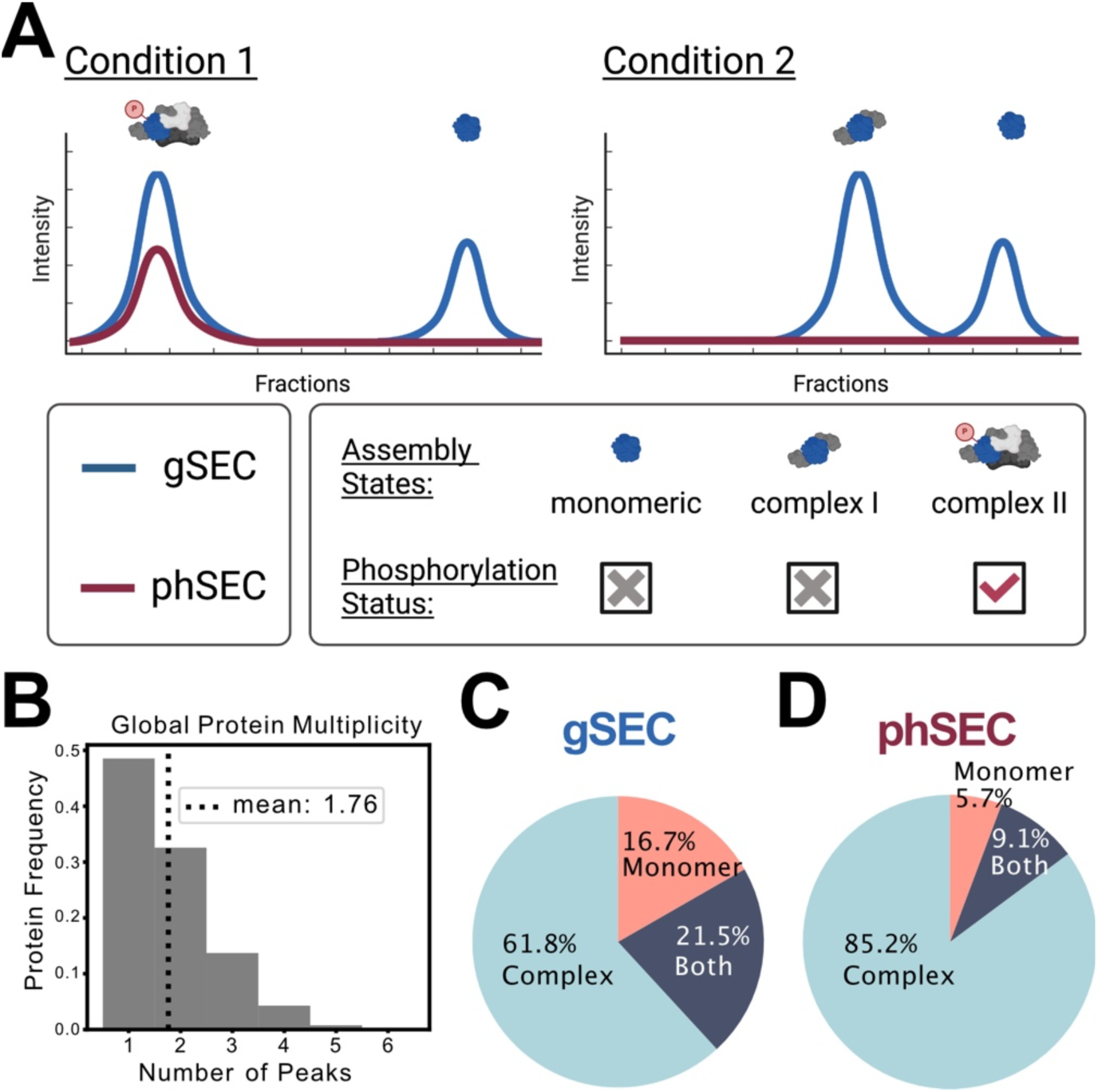
A novel protein-centric analysis framework to study assembly-states: **(A)** Hypothetical SEC-MX elution traces depicting a protein with multiple assembly-states, each represented as a distinct peak. Elution peaks can also differ between conditions; for example, under condition 1, the depicted protein elutes as a monomer (not phosphorylated) and a large complex including a phosphorylated-site. Under condition 2, the monomer peak is still evident, while the different position of the larger MW elution peak suggests that the protein is taking part in a different complex, which is not phosphorylated. **(B)** Frequency distribution of peak multiplicity per protein showing that >50% of proteins elute with >1 peaks. **(C-D)** Pie charts showing the percentage of proteins that eluted exclusively with a peak categorized as monomeric, complexed, or both (based on MW estimation) in gSEC (C) and phSEC (D).

The first step in our analysis pipeline was to define assembly-states per protein. We used a peak-calling algorithm (as detailed in Materials and Methods) to identify assembly-states for each protein in each dataset independently (gSEC/phSEC, for each cell-line and biological replicate). This analysis identified that the average ∼5000 proteins in each gSEC sample are eluting in ∼8,500 assembly-states, with an average multiplicity of 1.76 peaks per protein, and 1.4 peaks per peptide (Table 1, Figure 3B, Extended Data Figure 3F), suggesting that over half the proteins in the cell are present in at least two different assembly-states.

Following peak calling, each assembly-state was categorized as monomeric or complexed based on the peak position along the SEC dimension (detailed in Materials and Methods). We observed that only ∼17% of proteins eluted exclusively as monomeric, 62% exclusively in their complexed form, and 21% in both (Figure 3C), suggesting that the majority of proteins in the cell are complexed with other molecules, in line with previous observations ^1,2,15^. This trend toward assembled proteins was even more prominent for phSEC profiles, where less than 6% of proteins eluted exclusively as monomeric, 85% exclusively in their complexed form, and 9% in both (Figure 3D). Comparing these distributions, we observed that the phSEC assembly-states eluted less frequently as monomers than in gSEC, a finding that we confirmed for the subset of proteins matched with measured phosphopeptides (Extended Data Figure 3G). Moreover, the lower percentage of monomeric assembly-states in phSEC compared to gSEC suggests that phosphorylation occurs more often on the assembled form of a given protein, rather than its monomeric.

In conclusion, we used a peak-focused approach to define assembly-states from SEC data. Using this method, we showed that more than half of the measured proteins presented in two or more assembly-states, highlighting the importance of considering how alternative assembly-states might be regulated. Therefore, we next turned to assign phosphorylation-events to specific assembly-states.

### Mapping PTM onto assembly-states

To map phosphorylation onto assembly-states, we aligned the peaks identified in phSEC to the peaks of their parent-protein in gSEC based on peak-apex position along the SEC range (see Materials and Methods). This analysis covered 1,592 proteins that were measured in both datasets, and showed that 90% of the phSEC proteins had at least one peak aligned between gSEC and phSEC, showing high agreement between the two datasets (1,446 out of 1,592, Figure 4A). When a phSEC peak aligned with gSEC peak, we considered that as evidence that the protein is phosphorylated in the specific assembly-state. Interestingly, many of the 1,446 proteins had gSEC peaks that were not aligned with the phSEC peak, suggesting that a substantial fraction of the phosphorylated proteins were selectively phosphorylated on specific assembly-states. Since this pattern can be affected by the lower coverage in the phSEC versus gSEC dataset, we filtered the proteins based on replicate reproducibility and delineated 257 candidate proteins for which phosphorylation differed between different assembly-states of the same protein (see Supplementary Material). We next discuss a few of these examples.

**Figure 4.**
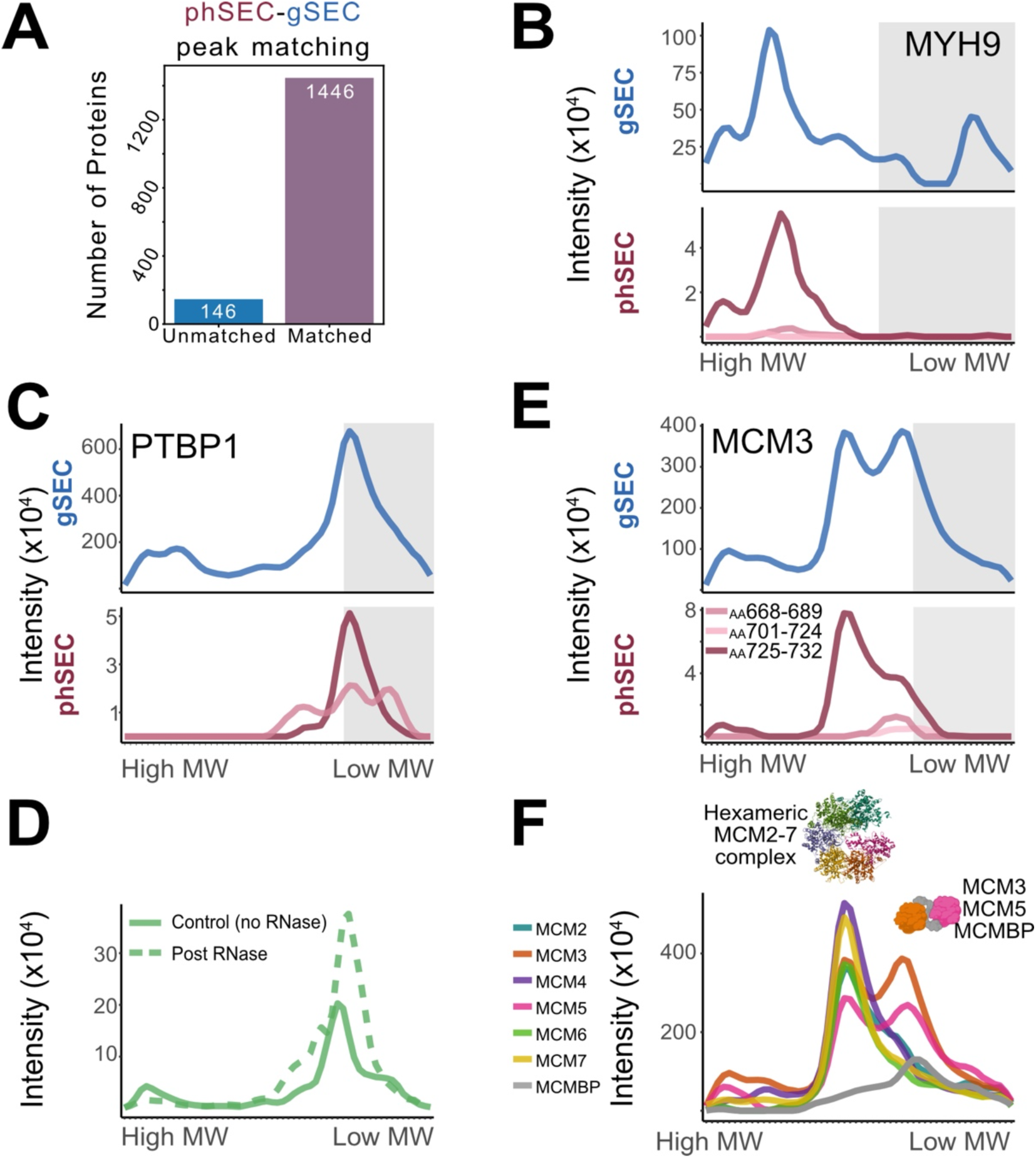
Mapping PTMs onto assembly-states: **(A)** Number of proteins for which phSEC peaks were matched to the gSEC peaks, based on their apex position: 1,446 out of the 1,593 proteins (90%) measured in both gSEC and phSEC had a least one phSEC peak that matched to a gSEC peak. **(B)** SEC-MX elution traces in gSEC (top) and phSEC (bottom) for MYH9 in HEK293 cells (averaged across replicates. HCT116 elution patterns shown in Extended Data Figure 4). Gray boxes indicate the range of fractions covering the monomeric form of the protein. **(C)** Same as B, for PTBP1. **(D)** PTBP1 elution in SEC-DIA performed on HEK293 cell lysates before/after RNAse treatment. **(E)** Same as B, for MCM3. **(F)** gSEC elution traces for all MCM2-7 complex subunits (HEK293, averaged across replicates) support the existence of two assembly-states: the full complex and an intermediate assembly-product comprising of MCM3-MCM5-MCMBP. Crystal structure of human single hexameric MCM2-7 complex is shown (PDB 7W68, deposited by Xu, N.N. et al., 2021-12-01).

Among the candidate proteins we found the non-muscle myosin IIA Myosin-9 (MYH9), an abundant actin-motor expressed in most eukaryotic cells. MYH9 eluted in both a high MW peak (multimeric assembly-state), and a smaller MW peak in the monomeric range (Figure 4B, Extended Data Figure 4A). Interestingly, phosphopeptides spanning the sequence around the known phosphorylation site on Serine-1943 were reproducibly measured in phSEC, all co-eluting only with the multimeric assembly-state, suggesting that MYH9 is differentially phosphorylated between its monomeric and assembled forms. This observation is in-line with previous works showing that association of MYH9 with its binding partners is regulated by Serine-1943 phosphorylation^31,32^.

In addition to multimeric assembly-states determined by PPIs, assembly-states may form by interaction with various other molecules, such as nucleic acids. One of the advantages of using our protein-centric approach for analyzing SEC data, as opposed to a PPI focused approach, is the ability to identify that a given protein has significant assembly-state changes without the requirement to identify the underlying interactors. For example, we observed that several RNA binding proteins (RBPs) in our data eluted in multiple gSEC peaks that seemed to be differentially matched with a peak in phSEC. For instance, Polypyrimidine tract-binding protein 1 (PTBP1) eluted in two prominent assembly-states; in fractions 8 (high MW) and 40 (low MW). Interestingly, PTBP1 phosphopeptides eluted within a single peak overlapping the smaller MW assembly-state (Figure 4C, Extended Data Figure 4B), suggesting that only this assembly-state of PTBP1 is phosphorylated. One potential explanation to this phenomenon could be that phosphorylation regulates the association of the protein to RNA transcripts. To test this hypothesis, we plotted the elution profile of PTBP1 in SEC from HEK293 cells, before and after RNAse digestion (Figure 4D). We observed that the intensity of the high MW peak of PTBP1 is decreased following RNAse digestion, and the smaller MW peak is increased – supporting the notion that the phosphorylated PTBP1 form is no longer interacting with RNA. A similar pattern was observed in our data for various other RBPs including CPSF3, which like PTBP1 seems to be phosphorylated only in its RNA-free form (Extended Data Figure 4C). On the other hand, other RBPs, like DDX54 (Extended Data Figure 4D), exhibited the opposite pattern with phSEC peaks only matching the high MW RNA-bound assembly-state. Together, these observations suggest a role for phosphorylation in regulating RBPs interactions with RNA.

Another layer of information in our matching analysis comes from comparing the patterns of different phosphopeptides mapped to the same parent-protein. In some cases, although all assembly-states were identified as phosphorylated, a deeper dive in the data showed that each assembly-state is phosphorylated on a different site, as in the example of the DNA replication licensing factor MCM3 (MCM3). MCM3 is a subunit of the minichromosome maintenance 2–7 complex (MCM2-7), a hetero-hexameric complex that functions as a DNA replication licensing factor. It is loaded onto replication origins to form inactive pre-replicative complexes, which are then activated by kinases to form the active CDC45-GINS-MCM helicase complex^33,34^. In gSEC, MCM3 eluted in two assembly-states: the full hexameric complex in fraction 24 (higher MW) and a lower MW complex in fraction 36 with MCM5 and the auxiliary protein MCM binding-protein (MCMBP), (Figure 4E-F, Extended Data Figure 4F). In phSEC, we identified three distinct phosphopeptides of MCM3, each containing known phosphorylation sites; one spanning AA 668-689 including Serine-672 and Serine-681, the other spanning AA 701-724 including Tyrosine-708 and Threonine-722, and the third spanning AA 725-732 including Serine-728. Interestingly, while phPEP^725-732^ co-eluted with both gSEC peaks, phPEP^701-724^ and phPEP^668-689^ eluted only in the lower MW peak (Figure 4E, Extended Data Figure 4E). Of note, a recent study suggested that MCMBP may play a role in forming the MCM2-7 hexameric complex before it is loaded onto the chromatin^35^, meaning that the MCM3-MCM5-MCMBP complex is potentially a stable intermediate assembly product of the full complex. Therefore, the observation that phPEP^668-689^ and phPEP^701-724^ elute only with the MCM3-MCM5-MCMBP assembly-state raises the hypothesis that the sites within these peptides are dephosphorylated prior to the assembly of the full complex. While further studies are required to test this hypothesis, these observations exemplify the potential of our method in formulating testable hypotheses of how phosphorylation contributes to differential assembly-states and functions.

In conclusion, we used a peak-focused analysis to map phosphorylation events to individual assembly-states. We showed how this approach can be used to explore the relationship between protein-interactions and PTM regulation. Furthermore, we found hundreds of potential cases of differential phosphorylation between assembly-states, such as in cases where the monomeric and assembled forms differ in phosphorylation (MYH9), how phosphorylation may regulate the binding of proteins to nucleic acids (PTBP1, CPSF3, DDX54), or the assembly-process of large multimeric complexes (MCM3). Next, we expanded the analysis to compare assembly-states and their phosphorylation between the two analyzed cell-lines.

### SEC-MX enables differential analysis between biological samples

One of the main considerations in designing SEC-MX was to facilitate differential analyses of assembly-state changes between different biological samples and/or conditions. In this study, we focused on analyzing the assembly-state changes and their phosphorylation-state between two human cell-lines, HEK293 and HCT116, whose PPIs were previously compared^36^. We analyzed differences between the samples at the peak level as most proteins in our study were observed eluting in more than one assembly-state. We speculated that a peak-level approach would be highly sensitive to the detection of proteins differentially regulated in only one of several assembly-states. Overall, the SEC profiles of HEK293 and HCT116 correlated very well in both gSEC and phSEC (Extended Data Figure 5A), with ∼70% of gSEC assembly-states and 63% of phSEC assembly-states identified in both cell-lines (Figure 5A-B). In accordance with previous studies, these observations indicate substantial differences between the interactomes of the two cell lines, with 30-40% uniquely identified assembly-states in either of the cell-lines^36^. Of note, SEC-MX is well-suited for identifying condition-specific assembly-states since the mixing scheme provides nearly 100% overlap in peptide coverage between the biological samples. Meaning that, in the presence of a peak in one condition, absence from the other is most likely not due to disparities in coverage and can be more confidently interpreted as a difference in assembly-state. Therefore, we looked at the reproducible differences between HEK293 and HCT116 and identified 587 proteins with assembly-states exclusive to HEK293 and 672 exclusive to HCT116. Additionally, we found 76 proteins with assembly-states phosphorylated only in HEK293 and 111 only in HCT116 (Figure 5C, see Supplementary Materials for detailed lists).

**Figure 5.**
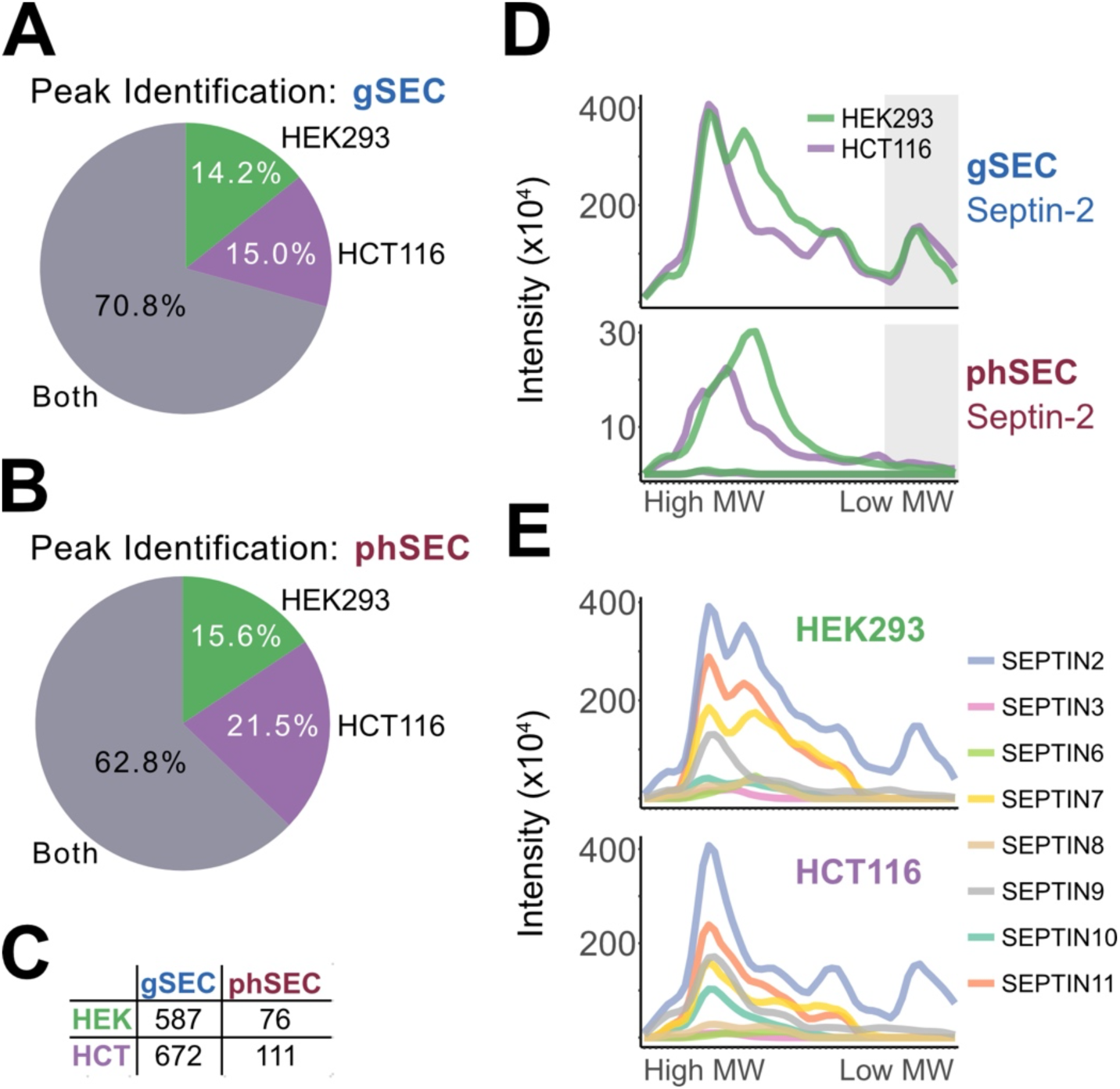
SEC-MX enables differential analysis between conditions: **(A-B)** Percentage of gSEC (A) and phSEC (B) peaks identified exclusively in HEK293 (green), HCT116 (purple) or in both (gray), across replicates. **(C)** Number of proteins with peaks identified exclusively in one of the cell-lines, reproducibly in both replicates. **(D)** SEC-MX elution traces for Septin-2, gSEC (top) or phSEC (bottom) in HEK293 (green) or HCT116 (purple). Gray boxes indicate the range of fractions covering the monomeric forms. **(E)** gSEC elution profiles for all measured Septin family members in HEK293 (top), or HCT116 (bottom).

Among the proteins with uniquely phosphorylated assembly-states we found the protein Septin-2 (Figure 5D). The Septin family is a group of conserved GTP-binding proteins, interacting with each other to form heteromeric complexes of 2, 3, or 4 members^37–40^. In HCT116 gSEC, Septin-2 eluted in one assembled form (fraction 11, high MW), and in its monomeric form (fraction 46, low MW). In HEK293 cells, an additional third assembly-state was identified in fraction 17 (Figure 5D). Overlaying the traces of all other identified Septins in gSEC, we observed that this additional assembly-state represents a different combination of Septins than the one in fraction 11, suggesting that different heteromeric complexes are formed in each cell-line (Figure 5E). Interestingly, the Septin-2 HEK293 phSEC elution pattern shows that the HEK293-exclusive assembly-state is more highly phosphorylated in this sample than its common (higher MW) counterpart (Figure 5D).

While our approach showed much potential for finding unique assembly-states, we were also interested in comparing the assembly-states common between conditions. Therefore, we quantified the abundance-differences of specific assembly-states by calculating the gSEC peak-ratios between the two conditions (HCT116/HEK293). To support this quantification, we compared the gSEC peak ratios to the total HCT116/HEK293 expression level ratios calculated based on shotgun DIA proteomics of the corresponding unfractionated samples (UF). We measured 4,163 proteins that were common between gSEC and UF and we observed a high correlation of the HCT116/HEK293 ratios measured by SEC-MX versus UF-DIA (Extended Data Figure 5B-C). Furthermore, we used an absolute log2 > 1 cutoff on the HCT116/HEK293 ratios in gSEC or UF to delineate differential proteins using each method and observed a ∼40% overlap between the candidate lists (Extended Data Figure 5D). Overall, the comparison to UF expression levels supports the use of the peak-focused quantification, while showcasing its potential to find differentially expressed assembly states.

Next, we applied the peak ratio analysis to discover differences in phosphorylation of assembly states using the phSEC dataset. Of the 1,446 proteins previously included in the analysis based on phSEC-gSEC peak alignment, we found 1,215 proteins for which at least one phosphorylated assembly-state was identified in both HEK293 and HCT116 (1,918 assembly-states). We then quantified the abundance-differences of specific assembly-states, or their relative phosphorylation, by calculating the peak-ratios (HCT116/HEK293) in gSEC and phSEC, respectively (Figure 6A). We used a cutoff of 2-fold difference in peak ratio to classify differential assembly-states at either the gSEC or phSEC datasets. These cutoffs divided our data to 9 groups based on whether assembly-states differ between HEK293 and HCT116 at the global level and/or phosphorylation level (Figure 6A). We observed that 788 proteins had assembly-states which were not significantly changed at either level (gNS/phNS). However, we found 88 and 89 proteins that had assembly-states enriched in HCT116 and HEK293, respectively, both at the abundance-level (gSEC) and the phosphorylation-level (phSEC), (groups B: gHCT/phHCT and C: gHEK/phHEK). Interestingly, our analysis discovered 301 proteins in HCT116 and 207 in HEK293 for which there was an assembly-state upregulated at the phosphorylation level, without a significant change in assembly-state abundance (groups A: gNS/phHCT, and D: gNS/phHEK). Moving forward, we focused on the four groups portraying differential phosphorylation and either concomitant abundance changes (groups B, C) or no significant abundance changes (groups A, D) (genes listed in Supplementary Materials).

**Figure 6.**
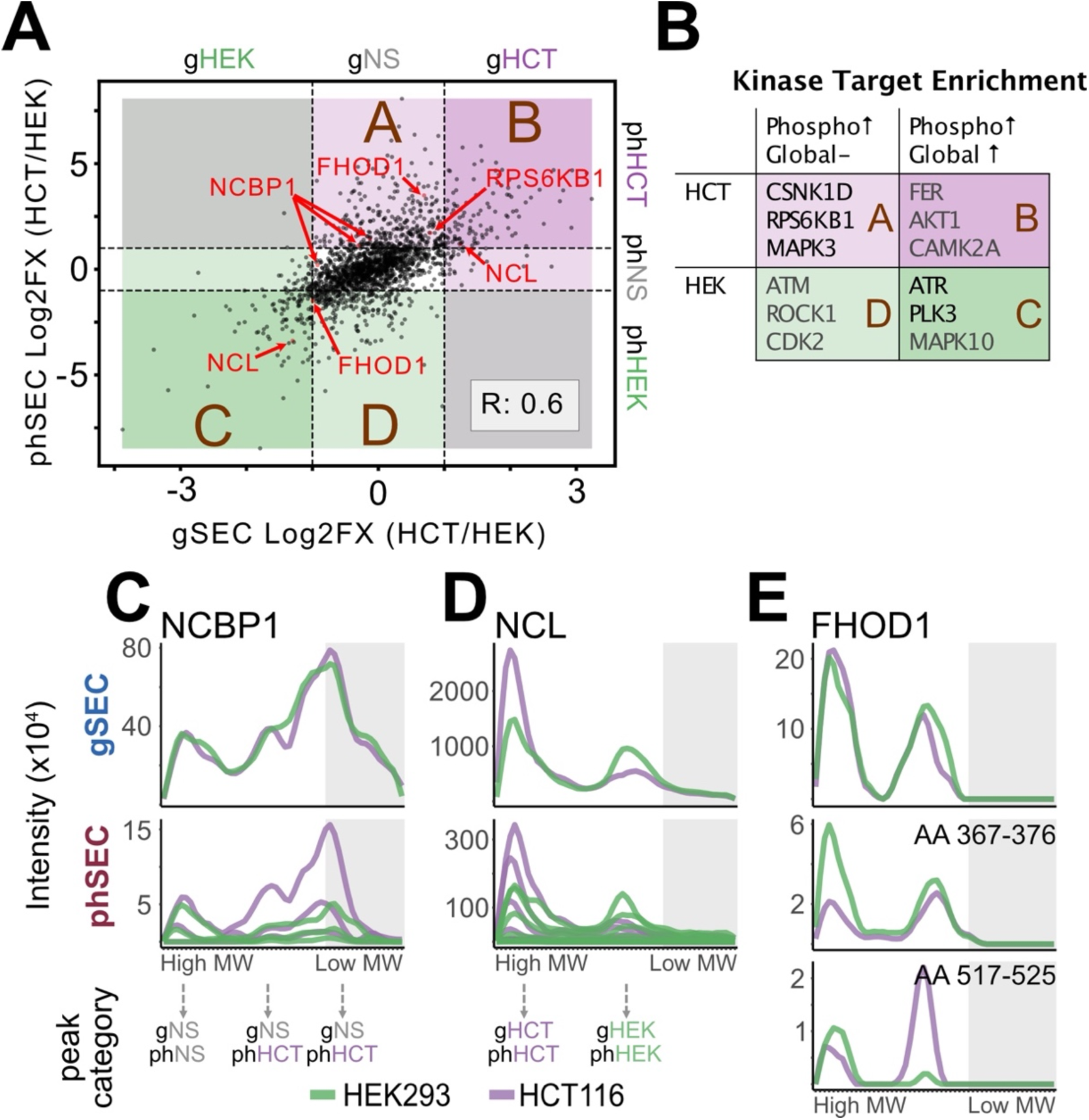
a peak-centric analysis uncovers different regulation of distinct assembly-states: **(A)** Scatter plot comparing the log2 ratio of peak heights (HCT116/HEK293) in phSEC (y-axis) and gSEC (x-axis), based on the subset of peaks identified in both cell-lines (gray area in Figure 5A-B). Dashed lines indicate the absolute log2 of 1, which was used as a cutoff for each dimension. These cutoffs divide the plot to 9 groups, based on whether assembly-states differ between the conditions at the global level and/or phosphorylation level, as indicated on the top and right sides of the scatter. **(B)** Kinase prediction by kinase target enrichment analysis results for each group indicated in A. Top 3 kinases are shown per group. Black font indicates q-value < 0.2 (FDR), while gray is q-value > 0.2. **(C-E)** SEC-MX elution traces for the indicated proteins (C-NCBP1, D-NCL, E-FHOD1), gSEC (top) or phSEC (bottom) in HEK293 (green) or HCT116 (purple). Gray areas indicate the range of fractions covering the monomeric form of each protein. For C-D, peak categories based on their position on the scatter (A) are shown on the bottom.

Given we identified 4 distinct groups, all enriched in phosphorylation (but not necessarily total protein changes – groups A and D), we hypothesized that each group may be regulated by different kinases. Therefore, we conducted a kinase target over-representation analysis^41,42^ on the proteins in each of the groups, which identified different potential upstream kinases in each group (Figure 6B). For example, group A (gNS/phHCT) was enriched in targets of the Ribosomal protein S6 kinase beta-1 (RPS6KB1). In group A, RPS6KB1 itself was enriched, along with its phosphorylation targets EEF2K, EIF4B, IRS1, MAPT, MTOR, RPS6, and NCBP1. As an example, we plotted the elution profiles of NCBP1, confirming it elutes in several peaks with similar abundance and distribution between HEK293 and HCT116 in gSEC, but with differential peaks in phSEC (Figure 6C).

RPS6KB1 is known to act downstream of mTOR signaling, promoting protein-synthesis by phosphorylating EIF4B, EEF2K and RPS6^43–49^. In-line with the observations of increased phosphorylation of these targets in HCT116, we found that ribosomal subunit expression (Gene Ontology (molecular function) enrichment^41^ on group B: gHCT/phHCT) and mTORC1-mediated signaling and Eukaryotic translation (Reactome pathway enrichment^41^ analysis on groups A and B: phHCT) were increased in this cell-line (Extended Data Figure 5E-F). Altogether, these observations highlight the power of our method as a tool for differential analysis between biological conditions.

As our data shows, proteins often elute with multiple assembly-states. These distinct assembly-states may be differently regulated, making them fall into different regions of our scatterplot, despite all being associated with the same parent-protein. For example, we identified that the RNA binding protein Nucleolin (NCL) had one assembly state upregulated in HCT116 (group B) and another upregulated in HEK293 (group C), indicating an inversed expression-pattern of the different assembly states (Figure 6D). Overall, we found that as many as 236 proteins had multiple assembly-states that were differentially regulated at the phSEC level (Figure 6A). For example, the FH1/FH2 domain-containing protein 1 (FHOD1) eluted in two peaks, which were not different in abundance in gSEC. However, one assembly-state (higher MW) was more highly phosphorylated in HEK293 and the other was more highly phosphorylated in HCT116 (Figure 6E). Notably, a closer inspection of the phSEC data showed that this pattern comes from distinct phosphopeptides: the phosphopeptide enriched in HEK293 spanned AA 367-376 including the known site on Serine-367, and the phosphopeptide enriched in HCT116 spanned AA 517-525 including the known site on Serine-523. Kinase prediction analysis^50^ showed these phosphorylation sites are most likely phosphorylated by distinct kinase groups, with Serine-367 (upregulated in HEK293) most probably regulated by kinases from the Aurora kinase family (AURB, AURC) and Serine-523 (upregulated in HCT116) most probably regulated by a kinase from the CMGC group (P38B, P38A, GSKB). Together, these observations support a model whereby FHOD1 is alternatively phosphorylated in an assembly-state specific manner. Meaning that, regardless of overall similar abundance of the two assembly-states, they are likely differentially regulated via phosphorylation between the two cell lines. This illustrates that integrating assembly data with phosphorylation status not only offers valuable insights into how PTMs influence protein interactions, but also sheds light on how PTMs contribute to distinct regulation patterns on different assembly states across samples.

In conclusion, we used our peak-focused approach to analyze differences in assembly-states between biological conditions, both at the full-protein (gSEC) and at the phosphopeptide level (phSEC). Our results brought forward proteins with assembly states exclusively found in one of the cell lines, as well as proteins with differentially expressed assembly states – both on a global and phosphorylation-level. Furthermore, we delineated unique cases of differential phosphorylation between distinct assembly states of the same proteins. Further inspection of these candidates exemplified how this method can generate testable hypotheses on the regulatory connections between phosphorylation and assembly-states. Together, our results support the utility of SEC-MX and the peak-focused analysis approach in providing valuable information on the dynamics of assembly-state regulation (abundance and phosphorylation) between conditions, beyond what is detected by analysis of expression levels.

## Discussion

In this study we developed SEC-MX, a multiplexed SEC-MS method that enabled the mapping of PTMs to assembly-states and performing differential quantification between conditions. We showed that SEC-MX performs as well as state-of-the-art label free SEC-MS methods in terms of coverage and PPI identification, and in a fraction of the needed LC-MS/MS runs. More importantly, we showed that SEC-MX enabled phosphopeptide enrichment downstream of SEC – generating a unique, proof-of-concept, dataset comprising the SEC elution profiles of thousands of phosphopeptides and their corresponding parent-proteins. We displayed how a careful multiplexing design can position SEC-MX as especially well-suited for quantifying differences between samples. Additionally, we analyzed these datasets with a focus on peak-level comparisons to overlay PTM information on assembly-states and/or compare them between biological conditions.

Our dataset provides a unique and rare systematic view of the interplay between phosphorylation events and assembly-states, highlighting proteome dynamics beyond abundance changes. In-line with previous reports^1,2,15^, we confirmed that the large majority of the proteins in the cell are assembled with other molecules. Expanding on this notion, our analysis shows that over half of the measured-proteins eluted in multiple assembly-states, and that PTMs may coincide with only a subset of them, and can change between conditions. We further exemplified how our approach can generate testable hypotheses regarding the relationships between phosphorylation events and molecular complex-assembly (Figure 4), and how they may differ between conditions (Figures 5-6).

In developing our protein-centric analysis approach, we decided to deviate from state-of-the-art protein-correlation profiling (PCP) that aims to uncover pair-wise PPIs (Prince^51^, EPIC^24^, SECAT^22,52^). Inspired by complex-centric programs such as CCprofiler^15^ and PCprophet^30^, we defined assembly “states” based on distinct SEC elution peaks and focused on individual proteins – how they change between samples, similar to other protein-centric approaches^2^. Our analysis framework proved to be a simple and comprehensive tool to compare assembly-states between conditions while overlaying PTMs. Additionally, we identified assembly states for nearly every analyzed protein/peptide, as opposed to methods focused solely on the complex/PPI level, which often result in limited networks, covering only a fraction of all measured proteins. However, future applications of SEC-MX analyses should attempt to tie-in PPI networks to provide the composition of different assembly-states.

Similar to SEC-MX, other techniques apply a protein-centric approach to detect dynamic changes in the functional proteome, such as thermal proteome profiling (TPP^53,54^), covalent protein-painting (CPP^55^) and limited-proteolysis mass spectrometry (LiP-MS^3^). These methods are sensitive in detecting protein functional changes that can result from a wide-array of mechanisms, including and not limited to, molecular interactions and PTMs. However, they are limited in their ability to identify the specific molecular-mechanism underlying an observed change in a given protein, requiring prior information such as structure, and/or detailed biochemical follow-ups.

Moreover, co-fractionation analysis following treatment with non-specific phosphatases (phospho-DIFFRAC^28^) also took a protein-centric approach to identify proteins with altered SEC elution profiles post-treatment. Although phospho-DIFFRAC can provide some degree of causality on the effects of phosphorylation on assembly-states, it lacks direct measurements of phosphopeptides and cannot shed light on the specific modified sites. For example, both our study and phospho-DIFFRAC suggested that the assembly-state of MYH9 is regulated by phosphorylation. However, we also pinpointed the specific phosphorylated peptide, which contained a modification site known to affect interactions. While SEC-MX cannot determine causality in the relationships between PTMs and assembly-states, we showed how it can serve as a valuable hypothesis-generating tool, guiding the design of follow-up studies into the causal effects of PPIs and PTMs. Furthermore, SEC-MX can facilitate the collection of detailed time-course measurements to validate these hypotheses.

In its current form, SEC-MX depends on data-dependent-acquisition (DDA), due to the use of isobaric tags. As with other DDA methods, this potentially reduces the depth of coverage, which can result in missing data points. Therefore, special attention should be made in the analysis step to make certain that intensities were at least measured in a given TMT-mix before claiming that a peptide does not have a peak in a certain region. To mitigate these issues, future development of SEC-MX should test the use of isotopic tags compatible with DIA methods, such as mTRAQ^56^, as they are expected to become available in >3 channels.

In conclusion, SEC-MX represents a significant advancement in the study of protein regulation and cellular functions by providing a comprehensive platform for the simultaneous characterization of post-translational modifications and assembly-states. Our method not only streamlines measurement processes and enables PTM enrichment but also offers a novel approach to explore the intricate interplay between these molecular events. By generating a proof-of-concept dataset encompassing the identification and quantification of assembly-states and PTMs across two distinct biological samples, we provide a granular perspective on protein regulation. Moreover, our focus on distinct assembly states highlights the diverse regulatory processes that proteins undergo, underscoring the necessity of novel analytical tools such as SEC-MX. While our study primarily examined phosphorylation for proof-of-concept, the versatility of SEC-MX allows for future exploration of additional modifications (e.g., acetylation), thereby broadening its applicability across various biological contexts. Ultimately, SEC-MX enables researchers to explore dynamic cellular processes and sheds light on the intricate mechanisms driving protein complexity and regulation.

## Materials and Methods

### Cell culture

HEK293XT cells (Takara Bio Lenti-X 293T, #632180) were provided by the Yeo lab at UC San Diego (SEC-DIA versus SEC-MX experiments), or purchased from ATCC (ATCC, CRL-3216, in HEK293 versus HCT116 experiments), HCT116 cells were provided by the Prives lab at Columbia University. HEK293 cells were cultured in DMEM (containing L-glutamine and Sodium pyruvate) and HCT116 in McCoy’s 5A media. Both media were supplemented with 10% Fetal Bovine Serum and Penicillin (100 U/mL) Streptomycin (100 μg/mL). Cells were grown to 80-90% confluency and were harvested at passages 6-20.

### Sample preparation for SEC

SEC sample preparation was as previously described in Bludau et al. 2020. Cells (25-40 million per sample) were harvested by scraping in ice cold PBS, washed and pelleted. Pellets were flash frozen in liquid nitrogen and stored in -80 °C. Upon thawing, cell pellets were lysed in cold lysis buffer (for TMT versus DIA comparisons: 150mM NaCl, 50mM Tris pH 7.5, 1% IGPAL-CA-630, 5% Glycerol; for HEK-HCT experiments: 50 mM HEPES pH 7.5, 150 mM NaCl, 0.5% NP40) supplemented with 50mM NaF, 2mM Na_3_VO_4_, 1mM PMSF, and 1X protease inhibitor cocktail (Sigma), followed by 10-30 minutes incubation on ice with intermittent vortexing. Cell lysates was then pre-cleared by 10 minutes centrifugation at 10,000g (4 °C) followed by 20 minutes of ultracentrifugation at 100,000g, 4 °C. To dilute detergents in the buffer, samples underwent buffer exchange on Amicon® ultra-0.5 centrifugal filter with 30 kDa molecular weight cutoff (Sigma) into 50 mM HEPES pH 7.5, 150 mM NaCl and 50 mM NaF in iterative steps of no larger than 1:3 dilutions. The final dilution ratio of the original lysis buffer to detergent free buffer was 1:50. The cell lysate was further cleared by 5 minutes of centrifugation at 17,000g, 4 °C. The concentration of the supernatant was measured by Nanodrop spectrophotometer (Thermo Scientific) and adjusted to 20-50mg/ml. Two mg of lysate were loaded on the SEC column per run.

### Size Exclusion Chromatography

Size exclusion was conducted on an Agilent 1260 Infinity II system operated with Agilent OpenLAB ChemStation software (version C.01.09). Two mg of cell lysate at 20-50mg/ml were loaded onto a Yarra SEC-4000 column (Phenomenex 00H-4514-K0, 3μm silica particles, 500A pores, column dimensions: 300 x 7.8mm) and fractionated in SEC running buffer (50 mM HEPES pH 7.5, 150 mM NaCl) at a flow rate of 1ml/min (first TMT experiment) or 0.5ml/min (all other experiments) and 100μL fractions were collected between minutes 6.5 to 16 or 11 to 30, respectively into 96 Well DeepWell Polypropylene Microplates (Thermo Scientific).

### Protein digestion and desalting

Following SEC fractionation protein concentration was measured using the Pierce™ BCA protein assay kit (Thermo Scientific) based on the manufacturer’s instructions. Equal volumes (∼80μL) from each of the fractions containing proteins (54 - 72) were subsequently processed. Proteins were denatured by incubation with an equal volume of urea buffer containing 8 M urea, 75 mM NaCl, 50 mM HEPES (pH 8.5) and 1 mM EDTA at 25 °C, 600 rpm for 20 mins in 96 Well DeepWell Polypropylene Microplates (Thermo Scientific). Proteins were then reduced with 5 mM DTT at 25 °C, 600 rpm for 45 minutes and then alkylated with 10 mM iodoacetamide (IAA) at 25 °C, 600 rpm for 45 mins in the dark. Proteins were then diluted in a ratio of 1:3 with 50 mM HEPES (pH 8.5) to lower the urea concentration less than 2M, and digested with trypsin enzyme (Promega) at 25 °C, 600 rpm overnight using 1:50 (enzyme: substrate) ratio. Digested peptides were acidified using formic acid and desalted on in-house packed C18 StageTips (two plugs) on top of 96 Well DeepWell Polypropylene Microplates as elaborated in Rappsilber et al., 2007 ^57^. For DIA measurements, dried peptides were reconstituted to a final concentration of 0.5μg/μL with 3% acetonitrile/ 0.2% formic acid. For TMT labeling purposes dried peptides were reconstituted in 50 mM HEPES (pH 8.5). An aliquot of 0.2mg of each non-fractionated sample (for HEK and HCT global protein expression analysis using DIA) was processed in a similar manner.

For the HEK-HCT dataset we used a direct labeling method. Following fraction selection based on BCA measurements, proteins were denatured by incubation at 95 °C, 600 rpm for 10 mins, followed by two cycles of 1 minute bath sonication. After samples cooled down to room temperature, proteins were reduced with 5 mM DTT at 25 °C, 600 rpm for 45 minutes and then alkylated with 10 mM iodoacetamide (IAA) at 25 °C, 600 rpm for 45 mins in the dark. Proteins were then diluted in a ratio of 1:3 with 50 mM 4-(2-hydroxyethyl)-1-piperazinepropanesulfonic acid (EPPS), pH 9.0. pH was adjusted to ∼ 8.2, and samples were subsequently digested with trypsin enzyme (Promega) at 25 °C, rpm 600 overnight using 1:50 (enzyme: substrate) ratio, and TMT labeled following digestion.

### TMT labeling

For SEC-MX experiments used to compare to SEC-DIA samples were digested and desalted as elaborated above. The resulting peptides were reconstituted with 50 mM HEPES (pH 8.5). For direct labeling peptides were labeled in the adjusted digestion buffer (50mM EPPS, pH adjusted to ∼8.2). This protocol resulted in a mean labeling efficiency of 98.3%, (comparable to the classic protocol, mean efficiency of 98.7%), while minimizing loss of material due to multiple desalting steps.

For all samples, peptides were labeled by addition of TMTpro™ 18 plex reagents (Thermo Scientific) into the sample at a ratio of 1:3 (peptide: TMT) by mass in a final volume of 29% acetonitrile. The labeling reaction was incubated at 25 °C, 600 rpm for 1 hour before being quenched with a final concentration of 0.3% hydroxylamine. Samples were then pooled as described in the pooling scheme and dried at least half of the volume to lower the acetonitrile concentration to less than 5%. The labeled peptides were then acidified using formic acid (pH <3) and desalted on C18 StageTips (two plugs), ^57^. The desalted peptides were dried and resuspended in 3% acetonitrile/ 0.2% formic acid for subsequent liquid chromatography-tandem mass spectrometry (LC-MS/MS) processing.

### Considerations for the choice of TMT multiplexing schemes

We initially attempted to perform IMAC enrichment on SEC fractions multiplexed using the “full overlap” scheme, resulting in a low number of recovered phosphopeptides (∼1000 across all mixes). However, IMAC enrichment from SEC-MX mixes that were multiplexed with the “no overlap” scheme resulted in ∼5 times phosphopeptide identifications. Interestingly, comparing protein coverage in global datasets mixed with the full-overlap versus no-overlap schemes showed that protein coverage with the no-overlap scheme was only 10% lower, despite the 2-fold difference in measurements (3 versus 6 runs, for no-overlap versus full-overlap, respectively). However, the no-overlap multiplexing scheme yielded only a third of the PPI identifications using SECAT (Extended Data Figure 1). Therefore, we concluded that while the “full-overlap” scheme is more suited for interaction analyses, the “no-overlap” scheme was preferable for subsequent enrichment of phosphorylated-peptides.

### Phosphopeptide enrichment using immobilized metal affinity chromatography (IMAC)

IMAC enrichment from pooled TMT labeled peptides was conducted following the protocol by Mertins and colleagues ^58^. Briefly, Ni-NTA Superflow Agarose beads (Qiagen, cat. no. 30410, 20 µl beads / 40 µl slurry per sample) were washed in water 3 times, then stripped from nickel by 30 minutes incubation with 100mM EDTA, washed 3 times in water, and incubated with 10mM iron (III) chloride in water for 30 minutes in room temperature. After 3 additional washes in water, iron coupled beads were resuspended in binding buffer (1:1:1 (vol/vol/vol) ratio of acetonitrile/methanol/0.01% (vol/vol) acetic acid) in a final ratio of 1:1:1:1 beads/acetonitrile/methanol/0.01% (vol/vol) acetic acid. In parallel, TMT labeled peptides (post pooling) were resuspended to 0.5 µg/µl (∼100 µg per sample, estimated based on BCA assay of the SEC fractions) in 80% (vol/vol) MeCN/0.1% (vol/vol) Trifluoroacetic acid (TFA). Eighty µl of bead slurry were added to the peptides solution and incubated for 30 minutes at room temperature. Following this binding step, supernatants were aspirated and the coupled beads were resuspended in 200 µl of 80% (vol/vol) acetonitrile /0.1% (vol/vol) TFA in order to be loaded on C18 stage tips for desalting. Two-plug C-18 stage-tips were conditioned twice with 100% Methanol, washed in 50% (vol/vol) acetonitrile in 0.1% (vol/vol) formic acid (FA), and equilibrated twice with 1% (vol/vol) FA. Then, the enriched beads were loaded onto the stage tips. Loaded beads were washed twice with 50 µl of 80% (vol/vol) acetonitrile/0.1% (vol/vol) TFA, then twice with 50 µl of 1% (vol/vol) FA. Phosphopeptides were trans-eluted from the beads to the C18 material by three iterations of 70 µl of agarose-bead elution buffer (192.5 mM monobasic potassium phosphate / 307.5 mM dibasic potassium phosphate). Stage tips were washed twice in 1% (vol/vol) FA, and peptides eluted using 60 µl of 60% (vol/vol) acetonitrile / 0.2% (vol/vol) FA. Eluted peptides were dried using a savant speedvac and reconstituted in 15 µl of 3% (vol/vol) acetonitrile / 0.2% (vol/vol) FA.

### LC-MS/MS

LC-MS/MS analysis was performed on a Q-Exactive HF. 5μL of total peptides (at 0.5 µg/µL) were analyzed on a Waters M-Class UPLC using a C18 Thermo EASY-Spray column (2um, 100A, 75um x 25cm, or 15cm) or IonOpticks Aurora ultimate column (1.7um, 75um x 25cm) coupled to a benchtop ThermoFisher Scientific Orbitrap Q Exactive HF mass spectrometer. Peptides were separated at a flow rate of 400 nL/min with the following gradients: 70 minutes (SEC-DIA), 160 minutes (DIA runs for unfractionated samples), or 150 minutes (SEC-MX), all including sample loading and column equilibration times. For DIA runs MS1 Spectra were measured with a resolution of 120,000, an AGC target of 5e^6^ and a mass range from 350 to 1650 m/z. 63 isolation windows of 20 m/z were measured at a resolution of 30,000, an AGC target of 3e^6^, normalized collision energies of 22.5, 25, 27.5, and a fixed first mass of 200 m/z. For DDA runs MS1 Spectra were measured with a resolution of 120,000, an AGC target of 3e^6^ and a mass range from 300 to 1800 m/z. Top12 MS2 spectra were acquired at a resolution of 60,000, an AGC target of 1e^5^, an isolation window of 0.8m/z, normalized collision energies of 27, and a fixed first mass of 110 m/z.

### Data analysis

### Searches

Proteomics raw data were analyzed using the directDIA method on SpectroNaut v16.0 for DIA runs or SpectroMine (3.2.220222.52329) for DDA runs (Biognosys). Reference proteome used was human UniProt database (Homo sapiens, UP000005640, downloaded on August 8^th^ 2023). Search parameters were set to BGS factory settings for TMTpro 18 channels, modified without automatic cross-run normalization or imputation for SEC runs. Cross run median normalization and global imputation were used for global expression analysis (HEK-HCT non fractionated samples). Peptide spectral matches (PSMs), peptides and protein group data were exported for subsequent analysis. For phSEC, raw files were searched similarly on SpectroMine (3.2.220222.52329) with an additional variable phospho(STY) modification, with PTM localization workflow.

### Signal Processing

The peptide intensities were spread out along ∼55 SEC fractions. Peptides were filtered by being proteotypic and non-decoy. Empty or NA measurements were converted to zeros ^19^, and a single uniprot ID was assigned to each peptide. In TMT experiments, peptide reporter intensity values were normalized to their respective MS1 peak intensity. In experiments conducted with the full overlap TMT mixing scheme, TMT batch effects were corrected based on the signal in the common fractions between any two adjacent mixes. A normalization factor was calculated by dividing the peptide fraction intensities of mix [n+1] by mix [n], then taking the median of all the peptides and the mean of all the overlapping fractions in common between the mixes. Mix [n+1] was then normalized to mix [n] by multiplying all intensities by the normalization factor. Lastly, the peptide intensities of overlapping fractions were averaged.

Then, for the HEK-HCT dataset, the intensities were normalized between conditions. To do this, the median intensities were calculated in each sample (ptm/replicate/condition) and then the medians were used to find a total intensity ratio between the samples, which was then used to normalize HCT intensities to HEK intensities. Lastly, for plotting and matching purposes, the intensities were smoothed using <u>scipy.signal.filtfilt</u> a linear digital two-way filter (b=[1.0/2]*2, a=1).

Two normalized datasets were generated as described above to create smoothed elution profiles. The first consisted of protein-level global intensities and peptide-level phospho-enriched intensities. This dataset was generated to analyze the assembly-states on a protein level given the higher amount of protein ID overlap. The second dataset was on the peptide level for both the global and the phospho-enriched intensities. This dataset was used for certain comparisons where the peptide overlap between gSEC and phSEC would more accurately control for sources of variance.

### SECAT

SECAT was used to identify previously reported protein interactions ^22,52^. Replicates were analyzed in the same run to leverage the predictive power of the classifier. SECAT analysis was conducted on the processed peptide level signal (as mentioned above) using the default (SECAT provided) positive and negative interaction networks for the training step, and a target database of STRING’s human interactions (9606.protein.links.v11.5) for the query step. The default SECAT parameters were set except for a ‘pi0_lambda’ of 0.4 0 0 0, an ‘ss_initial_fdr’ of 0.5 and ‘ss_iteration_fdr’ of 0.2 during the ‘learn’ step. Additionally, the ‘export_tables’ option of the SECAT ‘learn’ step was used to export tables for extracting the STRING target and learning interactions along with their scores. The HEK-HCT data was also quantified by setting HEK as the ‘control_condition’, and using a ‘maximum_interaciton_qvalue’ of 0.1 for the quantify and export steps.

The networks were obtained by setting a q-value cutoff of 0.05 on the exported network tables. However, we initially observed that only 34% of the interactions were mutual to both DIA and SEC-MX datasets, but found a large number of interactions unique at a q-value < 0.05 cutoff were very close to the cutoff in the other experimental setup (Extended Data Figure 3B). Therefore, we adjusted the cutoff to include any interaction with a q-value between 0.05 and 0.1 (in at least 3 out of 5 SECAT runs), if its q-value was lower than 0.05 in the other dataset. With this adjusted cutoff we observed that 54% of interactions were identified in both datasets, while 29% were unique to DIA and 17% unique to TMT (Extended Data Figure 2B).

### EPIC

The EPIC tool was used to identify high confidence interactions allowing the discovery of novel interactions ^24^. Replicates were analyzed in the same run to leverage the predictive power of the classifier. Peptides were collapsed to the protein level by adding the top three peptide intensities for each protein. Proteins that eluted in only one fraction were filtered out. Pairwise protein-protein similarities were then computed using the Pearson Correlation-Coefficent (with and without noise), Jaccard, Apex, Mutual Information, and Euclidean metrics respectively. A cutoff of 0.5 for the features was chosen prior to analysis by a Random Forest Classifier which was trained on reference complexes generated using CORUM, INTact, and GO human proteins. The classifier was trained using an 80/20 cross validation split to minimize variance across runs and maximize predictive capabilities. Finally, de-novo protein-protein interactions were found by querying the classifier and reporting every interaction above 50% confidence as an interaction. To further benchmark the classifier a precision-recall graph was generated by varying the confidence of the classifier and reporting the metrics, the intersection of the precision and recall occurs at 60% confidence. However, as we are trying to minimize false positives, we picked a higher confidence of 80% (as previously reported by Pourhaghighi et al. 2020) which has less but more precise interactions. Confidence score cutoffs were further adjusted as elaborated for the SECAT dataset, to include any interaction with an EPIC score between 0.6 and 0.8, if its score was higher than 0.8 in the other dataset (Extended Data Figure 2E).

### Molecular weight (MW) estimation and monomeric fraction cutoff calculation

Molecular size estimations per SEC fraction were performed using a standard (Biorad 1511901), which was injected and measured onto the SEC column at the start and end of each experimental day. The calibration standard’s fractions and log MW were input into <u>sklearn.linear_model.LinearRegression</u> to create a log-linear model and predict the MW of each fraction. In this regard, the monomer fractions were predicted by using the Uniprot determined MW to get an estimated fraction of elution for the monomer. Additionally, a MW multiplier of 1.5 was used to account for wide or slightly shifted elution peaks.

### Peak Picking and Matching

Using the smoothed elution profiles (as describe in *Signal Processing*), peaks were identified using <u>scipy.signal.find_peaks</u> (prominence=max(intensity)*0.05, distance=3, height=1000) to get the apex position (the identity of the peak) and <u>scipy.signal.peak_widths</u> to get other attributes such as the peak height. In downstream analysis, peaks are synonymous with apex position. Each protein/peptide was expanded into a list of one or more peaks. When no peaks were identified, the protein/peptide was dropped due to noise.

With peaks identified in each sample, we then matched them within protein/peptide (protein-centric) between ptm enrichment and conditions. Peaks were matched within protein/peptide between condition/ptm/replicate by finding the best path. In order to do this, we first started with the list of identified peaks for a given protein/peptide in a sample (condition/ptm/replicate) as follows: *L_total_* = [*L*_1_, *L*_2_, … , *L_n_*], where L_i_ is the list of peaks for the i^th^ sample, out of ‘n’ samples.

Then, to account for non-matching peaks between samples (i.e., the peaks are too far away and therefore are not identified in one or more sample), we added NaN to each list of peaks (L_i_). The operation was performed such that 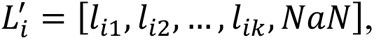 where l_ij_ are the peaks of L_i_, and ‘k’ is the final peak out ‘j’ peaks in sample ‘i’.

Next, we wanted to find all possible peak paths (Paths_all_) for a given protein *Path_all_* ={*Path*_1_, … *Path_n_paths_*}, where each path (Path_x_) is that protein’s alignment of peaks (*Path_x_* = {*Peak*_1_, … , *Peak_n_*}) between the different samples (1 to ‘n’). Furthermore, the total number of paths found for each protein/peptide can be expressed as the product of length of each peak list as follows 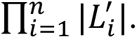 In other words, we generated all combinations of peaks (l_ij_) between each sample (L^’^_i_). Furthermore, each Path_x_ is comprised of a ‘n’ elements, one element for each sample (each element as either a peak or an NaN), where ‘m_peaks’ is the number of peaks in a Path_x_ (non-NaN values).

With all possible peak paths (Paths_all_), we wanted to find the best, non-redundant paths for each protein’s peaks to determine alignment. To do this, we first needed to score each path. First, we found the mean peak location of each path such that 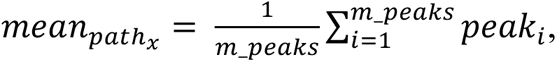 where peak_i_ is the peak (or NaN) selected in path_x_ for sample ‘i’. Then the paths were scored via a variance measure using a Threshold of 3 as follows:

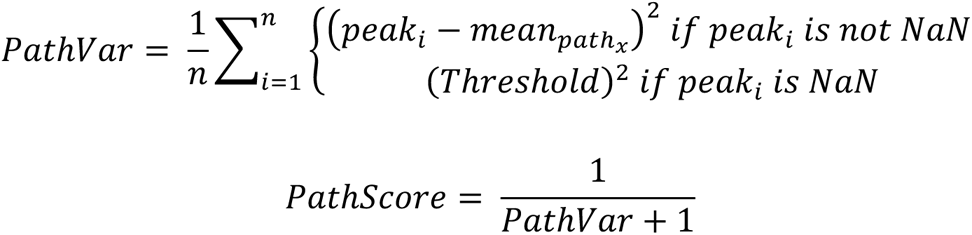

The Threshold was used to allow for the preferential selection of a given Path_x_ with an NaN in sample ‘i’ rather than a different Path where the variance introduced would exceed the threshold of 3 fractions. In other words, the Threshold value represents an estimate of the maximum permitted distance between peaks for matching.

With all the Paths (Paths_all_) each scored, we used an iterative method to select the lowest scoring 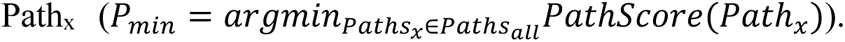 Then, we remove any other path elements from Paths_all_ that contain Peak_i_ from the P_min_ Path_x_, thereby making sure to only find the best non-redundant paths. Once the non-redundant paths are removed from Paths_all_, the lowest scoring Path_x_ is once again chosen (operation performed iteratively). This operation is iterated until all peaks are exhausted, whether matched with other peaks or by themselves:

Using the above formulas, the peak-level data was matched for each common protein between HEK and HCT116 as well as gSEC and phSEC. Fold changes were calculated as log2(HCT116/HEK293) for each assembly state based on identified peak height. When multiple phosphorylated peptides were identified for a given peak, their heights were summed before calculating fold changes. Additionally, peaks/assembly-states were classified as in the complex or monomer region based on the mean peak apex position of the matched peaks.

### Enrichment analysis

Enrichment analysis was performed using the WebGetalt (http://www.webgestalt.org) platform using the over-representation analysis (ORA) on Gene Ontology terms (molecular function non redundant) and Reactome Pathway^41,42^. Kinase target over representation analysis was conducted similarly on the same platform. Enriched sets were compared to a background list containing all the proteins identified in the experiment.

## Supporting information

Lists of proteins identified in various parts of the differential analysis

---

**Extended Data Figure 1.**
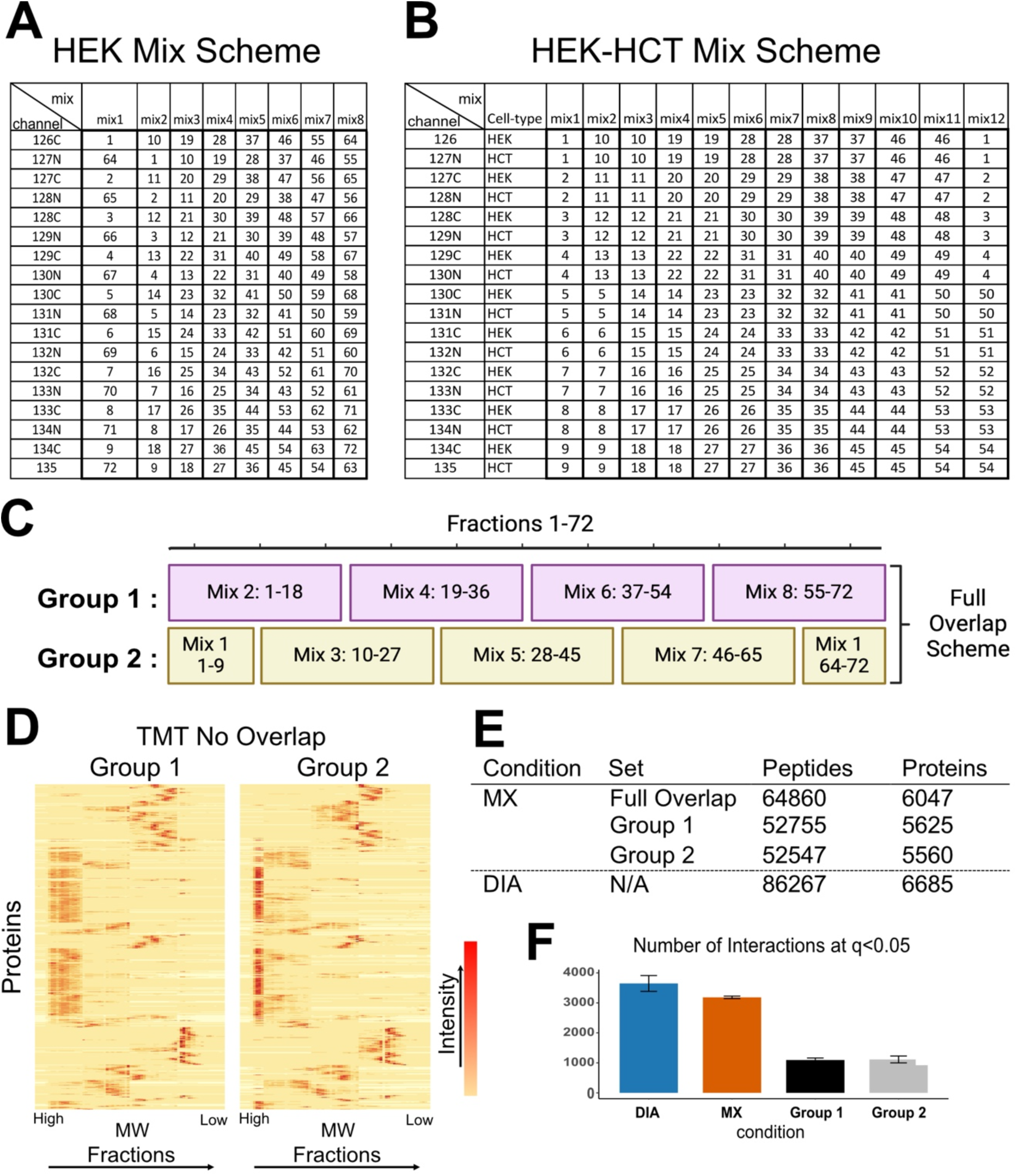
SEC-MX TMT Multiplexing: (**A**) Label used (TMTpro18) per fraction in the dataset comparing SEC-MX to SEC-DIA. (**B**) Label used (TMTpro18) per fraction in the datasets used for gSEC and phSEC from HEK293 and HCT116 cells. Replicate 1 was measured using a no-overlap scheme (only using the indicated mixes 1,3,5,7,9,11. Replicate 2 was measured with the full overlap scheme, as indicated (all 12 mixes). (**C**) A representation of the 72 collected fractions for the initial HEK TMT multiplexing. Group 1 consists of evenly numbered TMT mixes of all 72 fractions. Group 2 consists of the odd-numbered TMT mixes of all 72 fractions. The Full Overlap Scheme utilizes both TMT mixing Groups – with all fractions measured twice, and mixes from both groups in an adjacent staggered fashion to allow for inter-mix normalization. (**D**) Heatmaps of the protein intersect between both mix groups using the same dendrogram. The intensities are max-normalized to 1 for each protein. (**E**) Table of the numbers of peptides and proteins for both DIA and MX, with MX split between Group 1, Group 2, and Full Overlap. (**F**) Bar graph representation of the number of interactions identified with SECAT at a q-value less than 0.05 for DIA, MX (full overlap), Group 1 (MX), and Group 2 (MX).

**Extended Data Figure 2.**
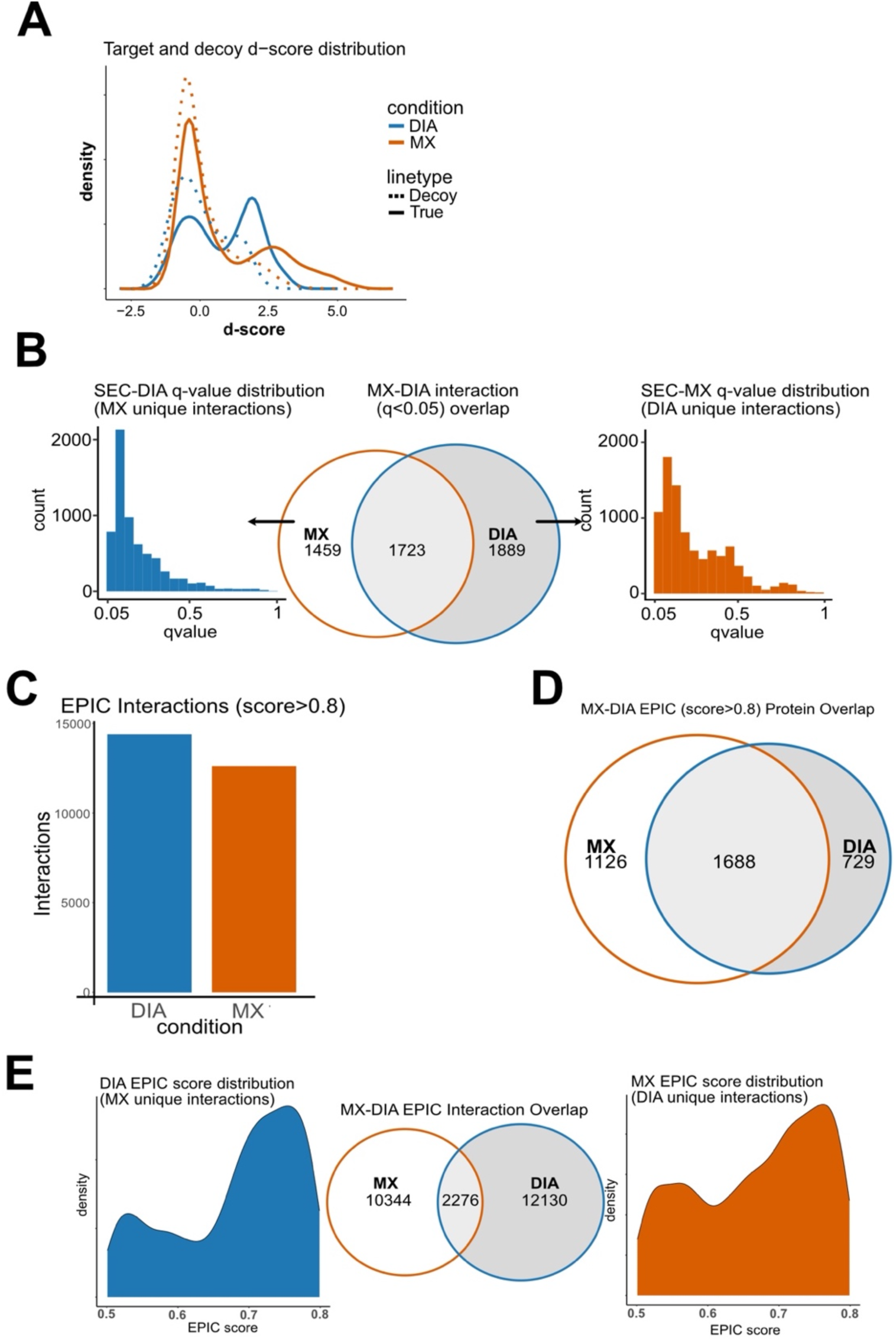
Interaction Analysis: (**A**) SECAT result of CORUM targets and decoys by discriminant score for both DIA and SEC-MX data. (**B**) Overlap of SECAT identified interactions at a q-value less than 0.05. For uniquely identified interactions, further q-value distribution plots for the opposite experiment are shown. (**C**) Barplot of the number of interactions identified using EPIC with a score cutoff of greater than 0.8. (**D**) Protein/node overlap between DIA and MX of the proteins identified in the EPIC interaction network. (**E**) Similar to (B), except for EPIC at a cutoff of 0.8.

**Extended Data Figure 3.**
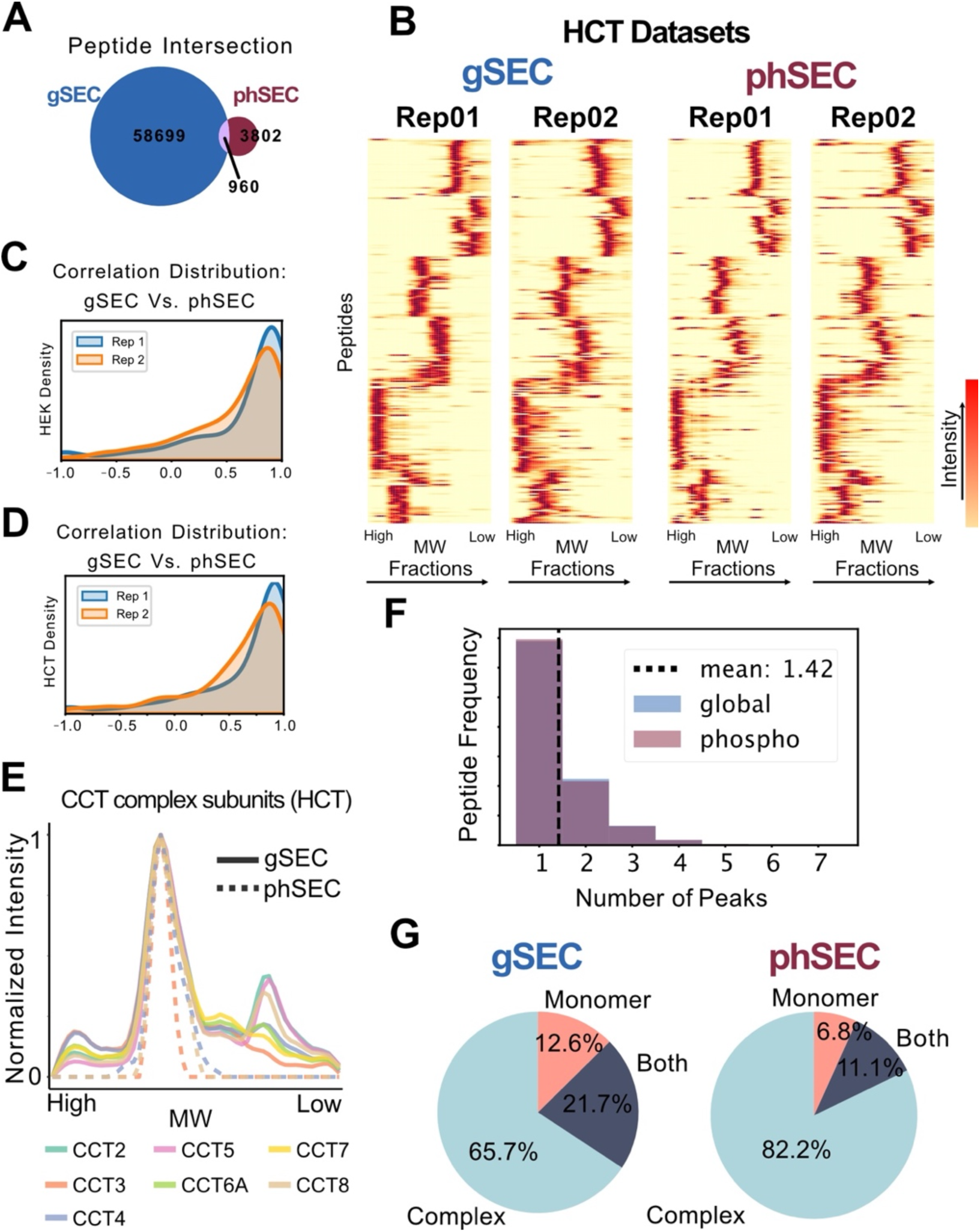
gSEC and phSEC datasets comparisons: (A) Venn diagram showing the overlap of peptides between the gSEC and phSEC datasets (based on stripped sequence). (**B**) Heatmaps of gSEC and phSEC by replicate for HCT116 data. Normalized to 1 by peptide maximum intensity and organized by the same dendrogram. (**C-D**) Distribution of Pearson correlation coefficients between gSEC and phSEC elution profiles for each overlapping peptide in HEK293 (C) and HCT116 (D), per replicate. (**E**) Elution profile plots for the CCT complex subunits in HCT116 (average intensity of replicates). Solid lines are from gSEC and dotted lines are from phSEC, and line color correspond to CCT subunit as shown in legend. (**F**) Stacked bar plot histogram of the number of peaks identified for each peptide intersecting in gSEC and phSEC (colors indicated in legend). Dotted line represents the mean number of peaks per peptide (same in global and phospho). (**G**) Pie-charts of protein peak distribution by fraction region (monomer, complex, or both) for gSEC (left) and phSEC (right) – for the intersecting proteins.

**Extended Data Figure 4.**
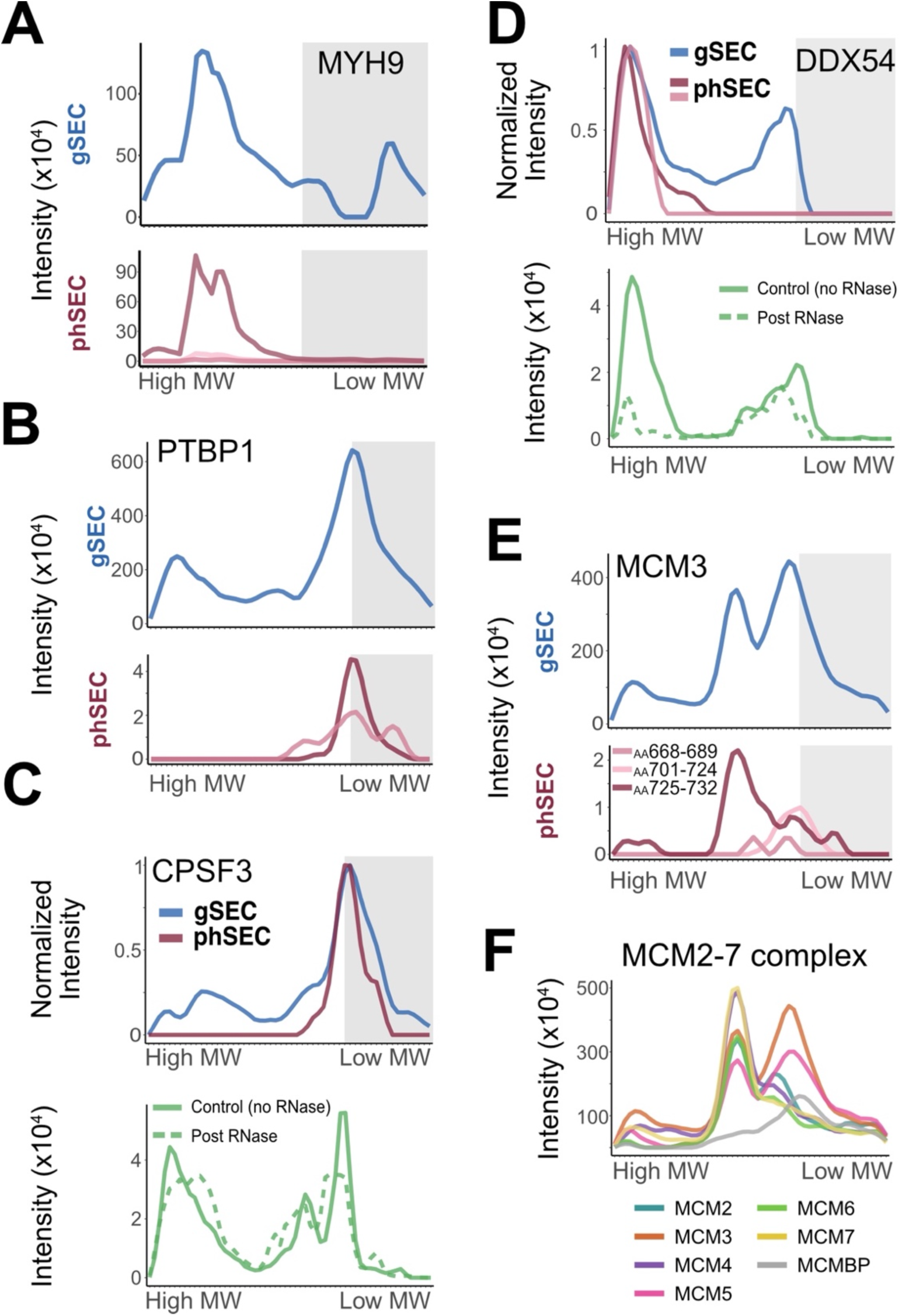
Additional Example Plots: (**A**) Replicate averaged elution profile plots of MYH9 for gSEC (top) and phSEC (bottom) in HCT116. Monomer region is grayed. (**B**) Similar plots for PTBP1 in HCT116 (see above). (**C**) On top is the normalized and replicate averaged elution profiles for CPSF3 in gSEC (blue) and phSEC (red) in HEK293. On bottom is the replicate averaged CPSF3 in SEC-DIA from HEK293 with RNase (dashed line) and without RNase (solid line). (**D**) Same as C, for DDX54. (**E**) Replicate averaged elution profile plots of MCM3 in gSEC (top) and phSEC (bottom) in HCT116. Monomer region is grayed. (**F**) Replicate averaged elution profiles for MCM2-7 complex subunits (see legend for subunits) in HCT116.

**Extended Data Figure 5.**
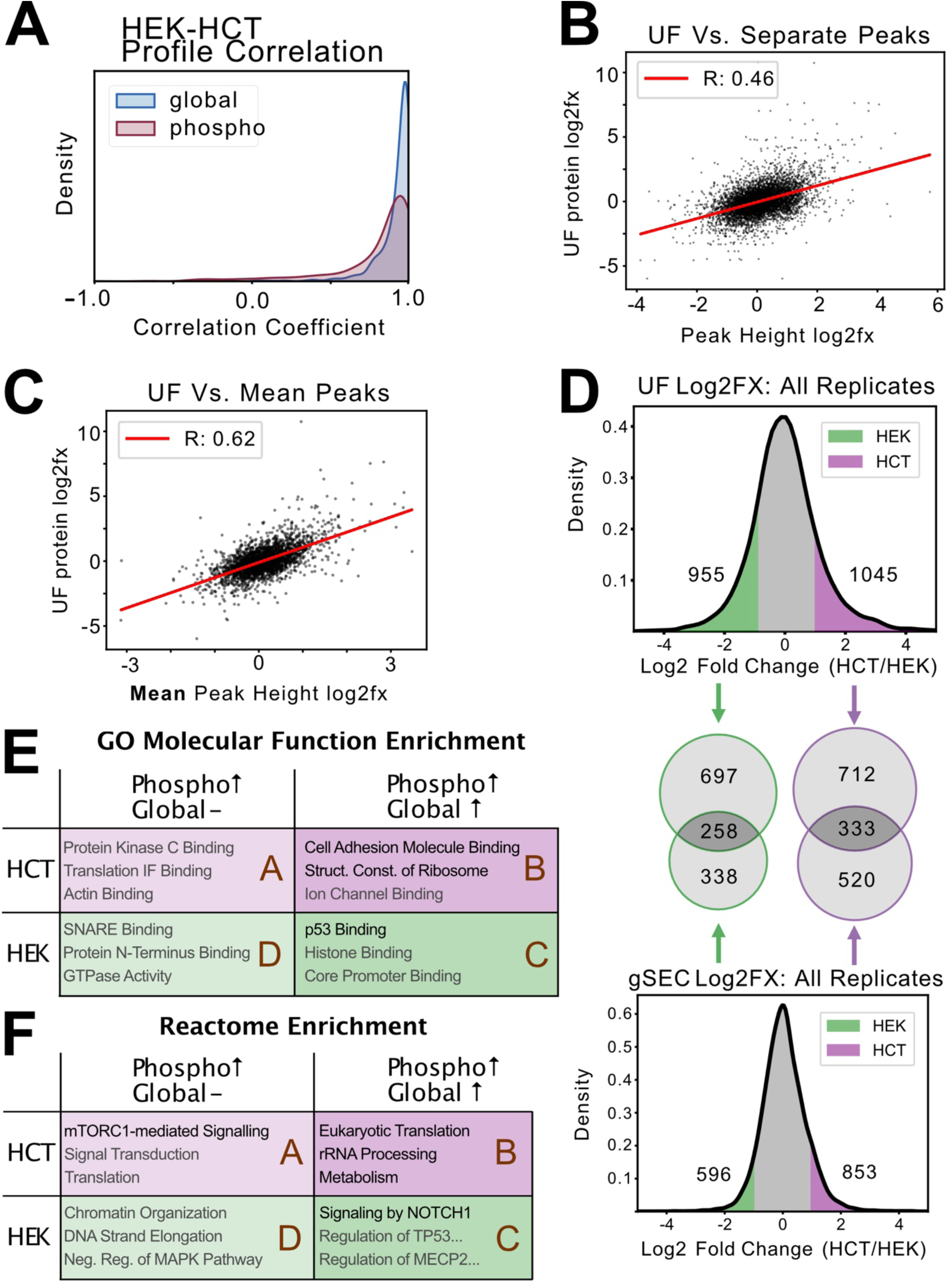
HEK-HCT Differential and Unfractionated Comparison: (**A**) Distribution of the Pearson correlation between HEK293 and HCT116 peptide elution profiles for the intersecting peptides in gSEC (blue) and phSEC (maroon). (**B**) Scatterplot of the log2 ratio (FX) between HCT116/HEK293 of the unfractionated analysis total intensity (y-axis) and the gSEC peak heights (x-axis). Regression line (red) is plotted with associated correlation (R=0.46). Multiple points for each protein’s peaks. (**C**) Similar to B, except the Peak Heights are averaged per protein. (**D**) Overall distribution of the Log2 Fold-Change with regions colored based on 2-fold increase cutoff for HEK293 (green) and HCT116 (purple). On top is the unfractionated total intensity distribution, and on bottom is the gSEC peak height distribution. Proteins found as significantly enriched are shown on the plots, with 955 and 1045 enriched in UF (HEK293 and HCT116 respectively), and 596 & 853 enriched in gSEC (HEK293 and HCT116 respectively). In between the Log2fx distributions, two Venn diagrams show the overlap in enriched proteins between UF and gSEC, with the left Venn for HEK293 (green), and the right Venn for HCT116 (purple). (**E**) Table of GO Molecular Function Enrichment for 4 different phSEC enrichment groups as determined by WebGestalt. Rows represent the cell-line of enrichment, and column represents whether a concomitant difference was observed in gSEC or not. Terms in black have an FDR < 0.2, and terms in gray have an FDR > 0.2. (**F**) Similar table as E, enriched for terms in the Reactome Pathway.

